# Benchmarking copy number aberrations inference tools using single-cell multi-omics datasets

**DOI:** 10.1101/2024.09.26.615284

**Authors:** Minfang Song, Shuai Ma, Gong Wang, Yukun Wang, Zhenzhen Yang, Bin Xie, Tongkun Guo, Xingxu Huang, Liye Zhang

## Abstract

Copy number aberrations (CNAs) are an important type of genomic variation which play a crucial role in the initiation and progression of cancer. With the explosion of single-cell RNA sequencing (scRNA-seq), several computational methods have been developed to infer CNAs from scRNA-seq studies. However, to date, no independent studies have comprehensively benchmarked their performance. Herein, we evaluated five state-of-the-art methods based on their performance in tumor vs normal cell classification, CNAs profile accuracy, tumor subclone inference and aneuploidy identification in non-malignant cells. Our results showed that Numbat outperformed others across most evaluation criteria, while CopyKAT excelled in scenarios when expression matrix alone was used as input. In specific tasks, SCEVAN showed the best performance in clonal breakpoint detection and Numbat showed high sensitivity in copy number neutral LOH (cnLOH) detection. Additionally, we investigated how referencing settings, inclusion of tumor microenvironment cells, tumor type, and tumor purity impact the performance of these tools. This study provides a valuable guideline for researchers in selecting the appropriate methods for their datasets.

## Background

Single-cell RNA sequencing (scRNA-seq) is a powerful and popular technology that provides a comprehensive view of the transcriptome of individual cells with high throughput and relatively low cost[1-3]. It has been successfully used to dissect the heterogeneous cellular tumor microenvironments, providing important insights into the biological functions of different cell subpopulations and cell-cell interactions, which are valuable for understanding cancer progression and therapy resistance[4, 5]. In addition to measuring gene expression levels, scRNA-seq can also reflect genomic aberrations such as copy number alterations (CNAs)[6-9]. Characterization of CNAs in tumor cells has significant implications for early tumor detection, delineating tumor heterogeneity, understanding modes of progression, and uncovering mechanisms of therapy resistance[10]. Single-cell DNA sequencing (scDNA-seq) is ideal for understanding CNAs at single-cell resolution [11-13]. However, the high cost and low coverage of scDNA-seq limit its application in estimation of copy number variations[14, 15]. To address this, several state-of the-art computational methods have been developed to infer CNAs from scRNA-seq data, enabling the delineation of intratumoral heterogeneity[16-20].

Current computational methods for inferring CNAs from scRNA-seq data can be broadly classified into two categories. One category is expression-based, where CNAs are inferred directly from gene expression. Examples of such tools include inferCNV[16], CopyKAT[17], and SCEVAN[18]. These tools operate on the principle that copy number amplifications (AMPs) or deletions (DELs) will lead to the upregulation or downregulation of genes within the affected genomic regions, respectively. InferCNV utilizes a corrected moving average over gene windows; CopyKAT employs an integrative Bayesian segmentation approach; SCEVAN uses a multi-channel segmentation algorithm. The second category combines gene expression with allele information, with relevant tools such as Numbat[19] and CaSpER[20]. Numbat employs a haplotype-aware Hidden Markov Model (HMM) and integrates signals form gene expression, allelic ratio derived from scRNA-seq, and population-derived haplotype to infer allele-specific CNAs. CaSpER utilizes a multiscale signal-processing framework that integrates gene expression and allelic shift signal profiles. Due to the differing principles, parameters, and use scenarios of these methods, it is crucial to evaluate their performance comprehensively.

In this study, we employed single-cell multi-omics datasets that enable simultaneous interrogation of DNA and RNA within the same cell. By using CNAs identified from scDNA-seq as the ground truth, we evaluated the accuracy of CNAs inference tools from scRNA-seq. Our comprehensive performance evaluation encompassed five popular tools: inferCNV, CopyKAT, SCEVAN, Numbat and CaSpER, from multiple perspectives. We examined the impact of reference settings, inclusion of tumor microenvironment (TME) cells, tumor type, and tumor purity on CNAs inference. Furthermore, we assessed the accuracy of tumor and normal cell classification, CNAs events accuracy, tumor subclone inference, and identification of aneuploid nonmalignant cells.

Our analyses suggest that Numbat demonstrated the best performance among the evaluated tools. In cases where only expression matrix is available, CopyKAT is recommended. In addition, our comprehensive study revealed common and specific pitfalls and their potential solutions. Therefore, we believe our benchmark study will serve as a valuable guidance for performing CNAs inference analyses from both single cell RNA-seq and spatial transcriptomics[21].

## Results

### Benchmarking framework and datasets examined

The benchmark dataset utilized in this study was collected from published studies that involved single cell multi-omics sequencing data, specifically paired scRNA-Seq and scDNA-Seq performed on the same cell. The CNAs inference obtained from scDNA-seq served as the ground truth. Due to the limited number of studies with paired scRNA-seq and scDNA-seq data, we selected samples with a sufficient number of cells from different sources. Specifically, we included eight colorectal cancer (CRC)[22] samples, two acute lymphoblastic leukemia (ALL)[23], one glioma, one neuroendocrine tumor and NPC43 cell line, along with HUVEC cell line samples[24] (**Table S1**).

In this study, we benchmarked five widely used tools for CNAs inference. The selection of these tools was based on two primary criteria: (1) they require only single-cell RNA sequencing (scRNA-Seq) data as input, and (2) they are recognized as popular tools within the field. (See Supplementary Note 1 for details). These tools can be broadly categorized into two major groups: those that solely require an expression matrix (inferCNV, CopyKAT, and SCEVAN), and those that require both expression matrix and B-allele frequency data (Numbat and CaSpER) **(Fig1**). The performance was assessed by three common applications in tumor cells: (1) classification of tumor and normal cells, (2) accuracy of CNAs events, (3) tumor subclone inference. Additionally, accumulating studies suggest that aneuploidy can also be present in normal cells, and it has been associated with various non-cancer diseases. Several publications have reported clonal expansion of CNAs in benign tissues[21, 25]. Furthermore, Zhou et al. identified prevalent CNAs in immune cells, endothelial cells, and fibroblasts within the TME of CRC[22]. Therefore, we also evaluated the performance of aneuploidy detection in normal cells.

**Figure 1:**
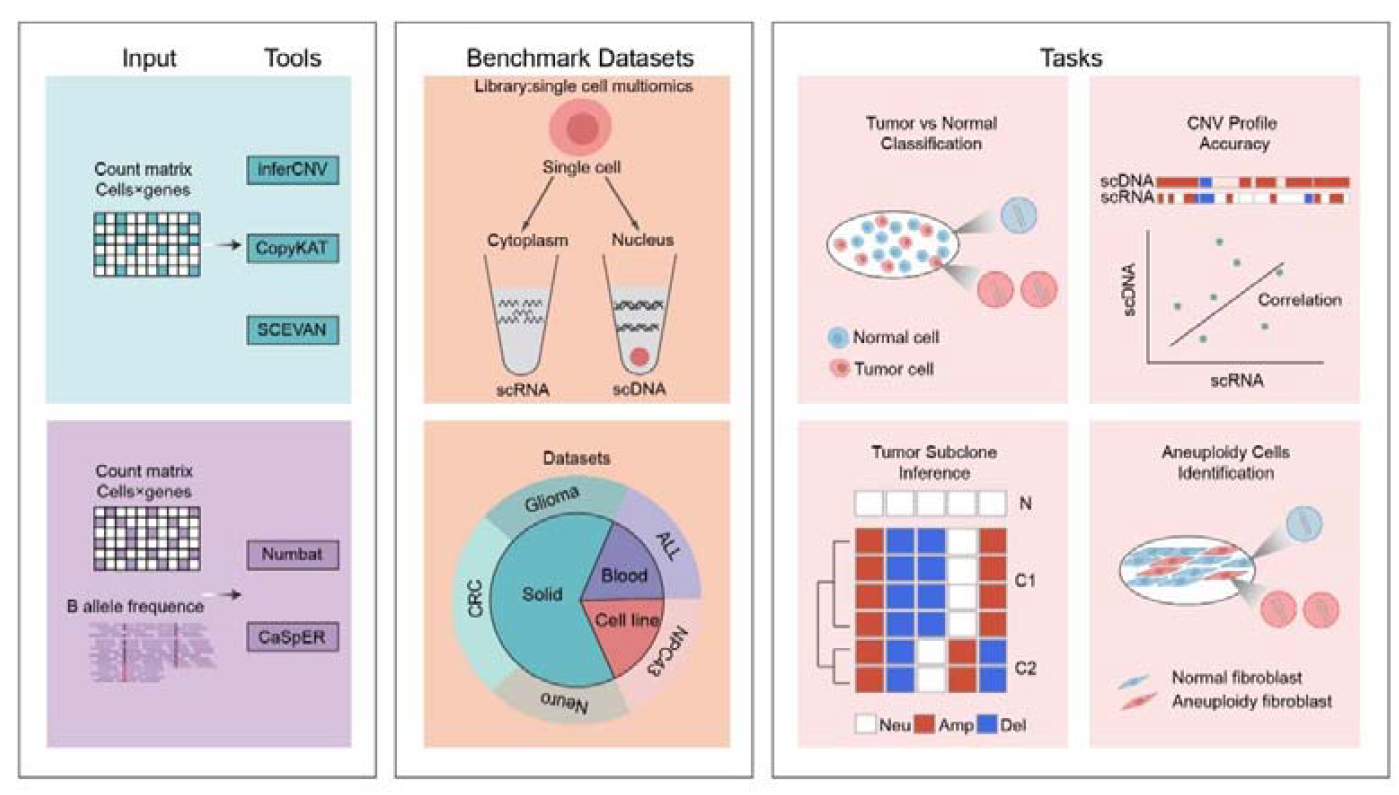
Overview of the benchmarking workflow. We evaluated five tools designed to infer copy number alterations (CNAs) from single-cell RNA-seq data. The methods fall into two categories: three rely solely on gene expression matrices: inferCNV, CopyKAT, and SCEVAN, while the remaining two, Numbat and CaSpER, utilize both gene expression and allele information. Our evaluation utilized single-cell multi-omics datasets that simultaneously interrogate DNA and RNA within the same cell, encompassing solid tumors, hematopoietic malignancies, and cancer cell lines. Copy number alterations (CNAs) detected from scDNA-seq served as the ground truth for evaluating the accuracy of CNAs inference tools based on scRNA-seq data. The performance of these five tools was assessed across several criteria, including tumor-normal classification, CNAs profile accuracy, subclonal structure inference, and aneuploidy cells identification in non-malignant cells.

### Distinguishing tumor cells vs. normal cells

To distinguish tumor cells and normal cells based on inferred CNAs profiles, we first applied the aforementioned tools to solid tumors including eight individuals with CRC, one patient with neuroendocrine tumor and one patient with glioma. Cell type annotations for all cells were obtained from corresponding published studies[22, 24, 26]. Only epithelial cells for CRC, endocrine cells for neuroendocrine tumor and glial cells for glioma were retained for tumor vs normal cell classification (**Figure 2A**). Overall, Numbat performed the best in distinguishing tumor cells from normal cells (**Figure 2B, C, Figure S1-S2A** and **Table S2**). Among the three tools that solely utilize the expression matrix, CopyKAT achieved the overall best performance. SCEVAN exhibited poor performance in samples with a low frequency of tumor cells (less than 40%), such as CRC11, CRC12, CRC16, and CRC15 (**Figure 2B and Figure S2A**). InferCNV was reliable except in CRC02, where the majority of cells are tumors (approximately 88%). In CRC02, the copy number gains and losses in tumors cells were incorrectly centered to be the copy number baseline by inferCNV, while normal cells were predicted to harbor the opposite CNAs profile and thus incorrectly classified to be tumor cells based on global copy number alteration levels (**Figure 2D, Figure S1F**). CaSpER also made the same error **(Figure S1F)**. To address this issue, we investigated whether including TME cells (such as immune, endothelial, and fibroblast cells) could improve the performance of inferCNV, CopyKAT, and SCEVAN, in cases with imbalanced tumor vs normal ratios (**Figure 2E, Figure S3A-C**). The inclusion of TME cells significantly improved the accuracy of tumor cell prediction for SCEVAN (**Figure2E, Figure S3C**). This improvement was reasonable because the SCEVAN algorithm incorporates a set of gene signatures from public collections, including cells from the tumor microenvironment such as stromal, immune cells, to identify high-confidence normal cells. Therefore, the addition of TME cells enhanced the ability to distinguish between tumor and normal cells for SCEVAN. Although this overall improvement was not observed in inferCNV, certain samples with a higher number of tumor cells, such as CRC02 (88%), CRC22 (91%), and CRC03 (67%), showed dramatic increases in their F1 scores, from 0, 0.5, and 0.55 to 1, 0.9, and 0.99, respectively (**Figure 2E, Figure S3A**). For instance, incorporating TME cells into the CRC02 sample not only improved the ability to distinguish between tumor and normal cells, but also accurately identified copy number alterations, including copy number amplifications on chr8 and chr20, and copy number deletions on chr15 (**Figure 2F**). This suggests that for inferCNV, when tumor purity is higher, including TME cells can enable more accurate estimation of expression level baselines, thus improving both tumor cell assignment and CNAs profile inference performance. This preliminary conclusion was further supported by the excellent performance of inferCNV in patient CRC15 where the tumor purity was exceptionally low (6%) (**Figure 2G**).

**Figure 2:**
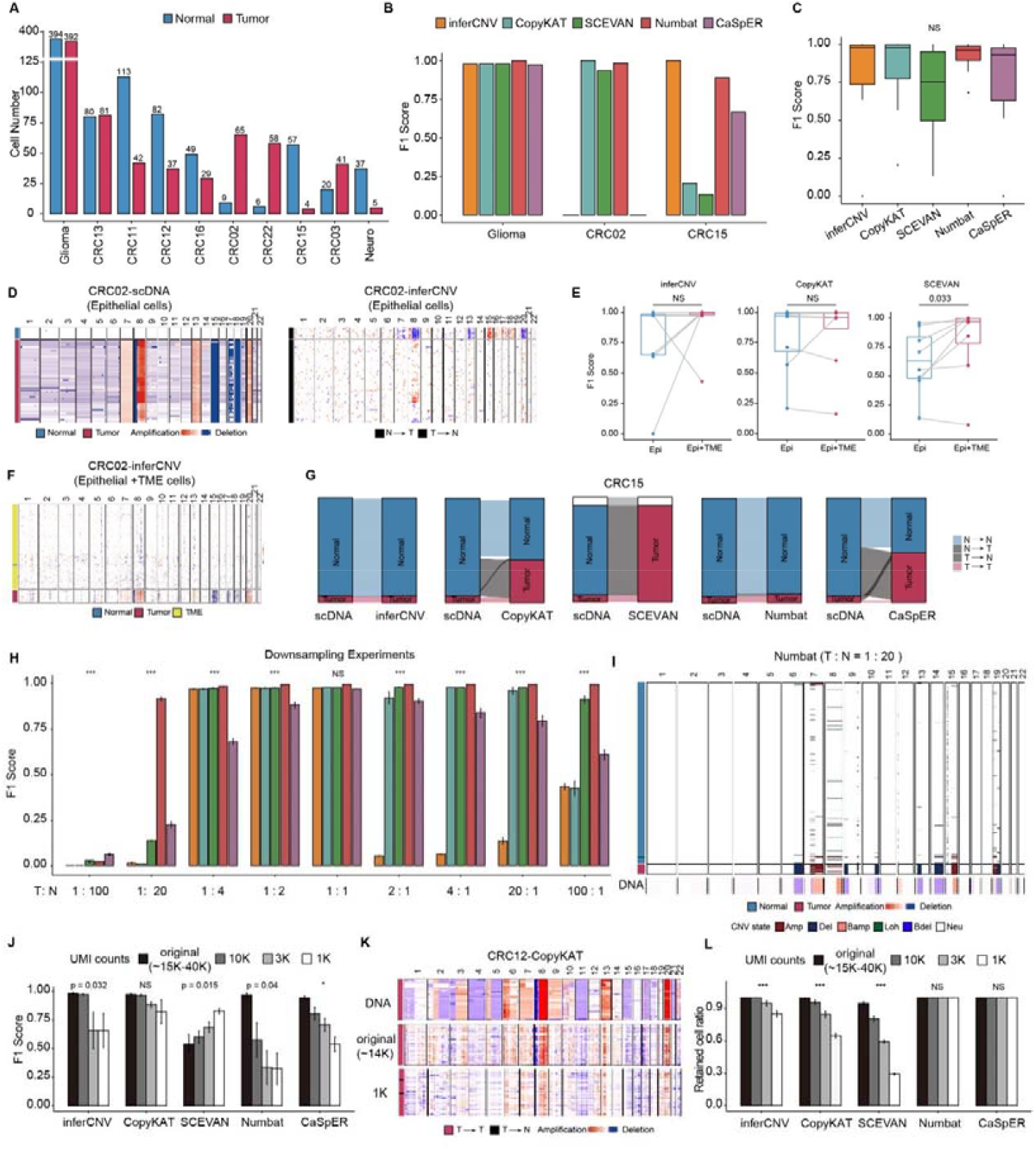
Benchmark of tumor and normal cells classification. A: A bar plot illustrates the counts of tumor and normal cells for each patient. B: The F1 scores for tumor and normal cell classification in Glioma, CRC02 and CRC15. C: Boxplot showed the overall F1 scores for tumor and normal cell classification in samples across the glioma, colorectal cancer, and neuroendocrine datasets. D: Heatmap showing the CNAs profiles of CRC02 epithelial cells derived from scDNA-seq (ground truth) (left side) and inferred from scRNA-seq by inferCNV (right). E: F1 scores between epithelial cells alone and epithelial cells combined with TME cells for CRC patients. F: Heatmap displaying the CNAs profiles of CRC02 of epithelial cells combined with TME cells estimated by inferCNV. G: Sankey diagrams illustrating the classification of tumor and normal cells from CRC15, with correct classifications as normal and tumor in both scDNA-seq and scRNA-seq data indicated by N>N and T->T, and misclassifications by N->T and T->N. T represents tumor cells, while N represents normal cells. H: F1 scores are shown for glioma downsampling experiments, with the x-axis indicating the ratio of tumor and normal cells. I: The upper heatmap represents the CNAs profiles of glioma from a downsampling experiment with a T : N ratio of 1 : 20, as inferred by Numbat. The lower heatmap represents the average CNA profile of tumor cells based on scDNA-Seq. J: F1 scores for tumor and normal cell classification under different sequencing depths. K: The heatmap demonstrates that CopyKAT less affected by sequencing depths. The upper heatmap represents the tumor cells CNAs profiles from scDNA-seq, the middle heatmap shows CNAs profiles from scRNA-seq, and the bottom heatmap exhibits CNAs profiles from down-sampled low-sequencing depth (1K). L: The ratios of cells passing quality control at different down-sampled sequencing depths. P values in E calculated using paired Wilcoxon rank-sum test. p values in (C, H, J and L) calculated using Kruskal-Wallis test. NS: p > 0.05, *: p < 0.01, **: p < 0.001, ***: p < 0.0001. T: Tumor cells, N: Normal cells. Additional data can be found in Figures S1-S5 and Tables S1-3.

To systematically explore the impact of tumor purity on the accuracy of CNA inference, we generated a synthetic dataset spanning a broad range of tumor/normal cell ratios (from 1:100 to 100:1) to examine the influence on the performance using the largest glioma dataset (**Figure 2H**). Numbat consistently outperformed the other tools, even in extreme cases with low tumor/normal cell ratios (**Figure 2I**). Notably, inferCNV exhibited opposite calling between tumor and normal cells when tumor purity was higher, consistent with the results observed in CRC02 (**Figure S4**).

To explore the impact of sequencing depth on the accuracy of CNAs inference, we’ve down-sampled the sequencing depth from several samples to a median of approximately 10k, 3k and 1k UMIs per cell. As sequencing depth decreased, the overall tumor-normal classification F1 scores for all tools dropped, especially in Numbat (Figure 2J and Figures S5A,D-E, Table S3). However, the performance of CopyKAT is largely unaffected. For instance, even at median of 1K UMIs per cell, CopyKAT could still correctly distinguish tumor from normal cells even though the CNA signal-to-noise ratios clearly decreased with the sequencing depth (Figure 2J-K and Figure S5A). The paradoxical increase in F1 scores in SCEVAN was mostly attributed to filtering out more nonmalignant cells due to lower sequencing depth, thus rectifying the prediction bias towards assigning most nonmalignant cells into malignant cells (Figure 2L and Figure S5A-D). It was noteworthy that cell filtering was implemented in inferCNV, CopyKAT and SCEVAN, but not in Numbat and CaSpER.

To conclude, in the cases with balanced malignant and non-malignant cells ratios, most tools performed equally well in tumor cell classification task, however, the performance of some tools deteriorated when the tumor vs normal ratios were very imbalanced. Numbat demonstrates the best performance when B allele frequency is available, while CopyKAT is recommended for cases where only the expression matrix is accessible or with lower sequencing depth.

### Accuracy of inferred CNAs profiles

To evaluate the accuracy of inferred CNAs profiles, it was imperative to ensure optimal performance of each software tool. In the process of inferring CNAs from scRNA-seq data, the designation of normal reference cells has a significant impact on the final CNAs results. Numbat and CaSpER software required the input of a reference, preferably derived from the same biological and technical conditions (**See method for more details**). For InferCNV, CopyKAT, and SCEVAN, the use of reference cells is optional. Given the potential problem of incorrect centering in inferCNV (as shown in Figure 2C) when using one-pass mode (single-run without specifying reference cells, **Figure 3A**), we hypothesized that employing a two-pass mode (first-pass run to identify normal cells and second-pass run using these normal cells as reference, **Figure 3A**) could improve the accuracy of CNAs profiles by facilitating baseline estimation. The similarity between CNAs profiles calling form scDNA-Seq and inferred CNAs profiles from scRNA-seq significantly increased in eight of nine cases for inferCNV (**Figure 3B-C**). In contrast, CopyKAT and SCEVAN already incorporated a two pass-like approach in their algorithms where they initially predict normal reference cells and then utilize them as a baseline to correct tumor cell CNAs[17, 18]. Thus, adopting a two-pass approach has minimal impact on the performance of these two methods (**Figure S6A-B**). Consequently, when assessing the accuracy of inferred CNAs profiles for inferCNV, the results obtained from the two-pass mode were utilized.

**Figure 3:**
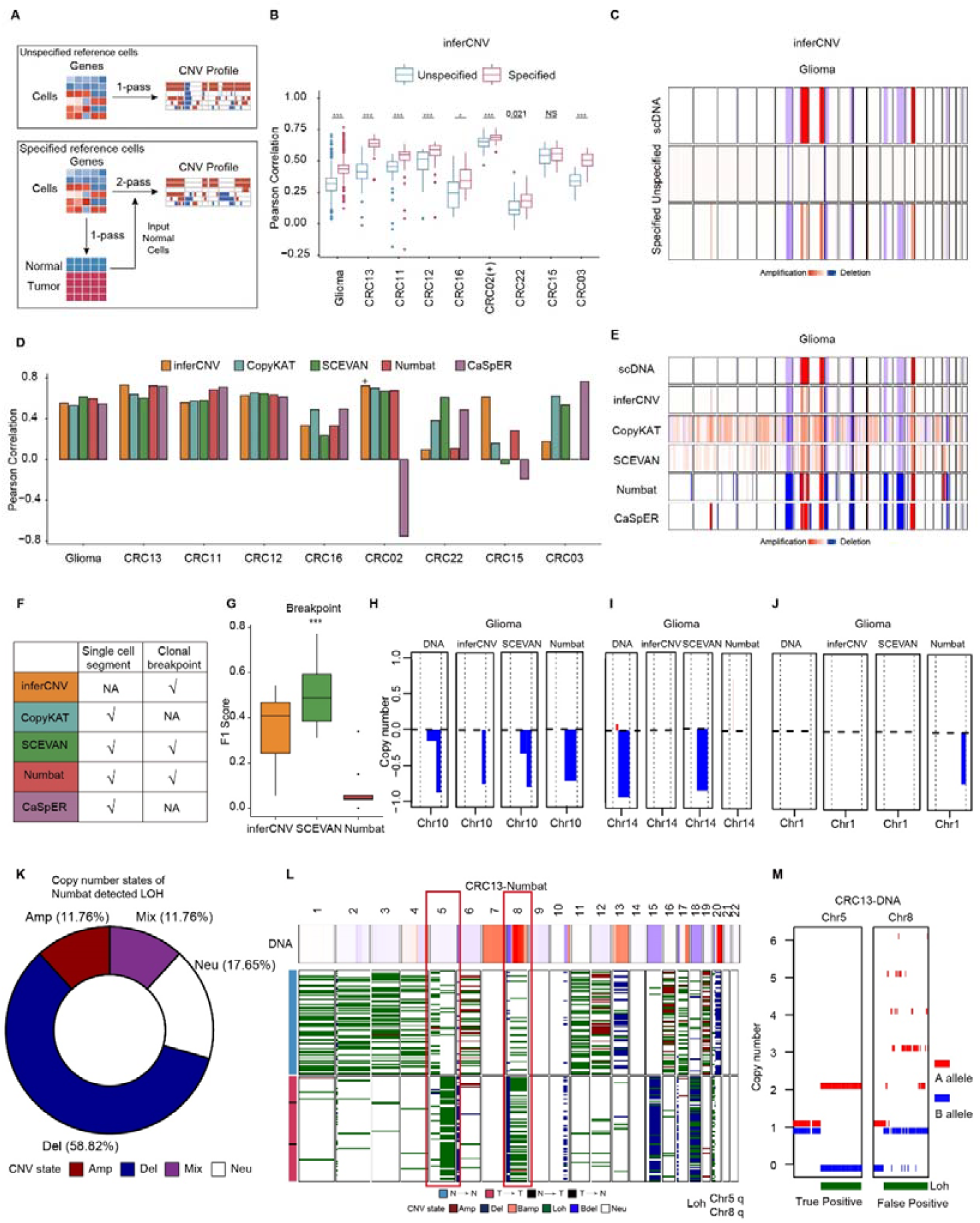
Benchmark of the accuracy of inferred CNAs profiles. A: Diagram illustrating CNAs profiles under unspecified (one-pass) and specified reference cells (two-pass) with inferCNV. In the one-pass scenario, inferCNV processes the whole cell population with the average as the reference. Conversely, in the two-pass approach, “normal” cells identified by inferCNV in the initial run (one-pass) serve as the reference for a second run (two-pass). B: Pearson correlation between inferred CNAs from scRNA-seq and the CNAs from scDNA-seq for each patient under unspecified (one-pass) and specified reference cells (two-pass) with inferCNV. Each dot represents the Pearson correlation between CNA profiles in scDNA-seq and inferred CNA profiles from paired scRNA-seq from the same cell. C: Heatmap showing CNAs profiles of glioma tumor cells obtained from scDNA-seq (upper), unspecified reference (middle), and specified reference using inferCNV (bottom). D: Bar plots illustrate the Pearson correlation of tumor cells between scRNA-seq inferred CNAs from and scDNA-seq for each patient across various tools. E: A heatmap displays the average CNAs profiles of glioma tumor cells derived from scDNA-seq assessed by different tools. F: Summary table for CNV segment and clonal breakpoint calling functionality among tools G: Boxplot exhibited F1 score for breakpoints in inferCNV, SCEVAN and Numbat H-J: Breakpoints from glioma samples in chromosome 10 (H), chromosome 14 (I) and chromosome 1 (J) K: The distribution of scDNA defined copy number status of cnLOH events detected by Numbat L: Heatmap showing the CNAs profiles of tumor/normal cells from Numbat in CRC13. Median copy number profiles from scDNA-Seq was shown on the top. Green regions are identified LOH regions. M: Sequenza CNA AB allele copy number results from combined scDNA-Seq on chr5 and chr8, respectively suggested that the LOH events were correct on chr5 and incorrect on chr8. “+” in (B) and (D) indicates tumor cell CNAs profiles estimated when both epithelial and TME cells are analyzed by inferCNV. p-values in (B) are calculated using unpaired two-tailed Wilcoxon rank-sum test and corrected using the Benjamini–Hochberg method. NS: p > 0.05, *: p < 0.01, **: p < 0.001, ***: p < 0.0001 Additional information is available in Figures S6-S11.

Numbat and CaSpER generated integral copy number profiles, which are easier to interpret and appear cleaner, while inferCNV, CopyKAT and SCEVAN output continuous copy number. No single tool consistently outperformed others in term of consistency between scRNA-Seq inferred profile and scDNA-Seq profile in all cases (**Figure 3D-E**). All tools performed well in cases with sufficient number of both tumor and normal cells, such as glioma, CRC13, CRC11 and CRC12. In samples with an imbalanced number of tumor and normal cells, such as CRC02, CRC22 and CRC15, the overall accuracy of CNAs profiles predicted by the five tools decreased considerably. Notably, in CRC03, although Numbat correctly assigned tumor cells, it accurately identified only a small fraction of deletion CNAs in high CNAs burden tumor cells, resulting in low similarity with CNAs profiles obtained from scDNA-Seq (**Figure 3D, Figure S1I**). We also evaluated the performance on hematopoietic cancer acute lymphoblastic leukemia (ALL). However, due to the unavailability of raw fastq data, we could only evaluate the performance of inferCNV, CopyKAT and SCEVAN (**Figure S7**). Among them, CopyKAT achieved the best performance, although noticeable false positive signals on CNAs were observed for both CopyKAT and SCEVAN (**Figure S7C, S7F**). Furthermore, we examined the performance of these tools on cancer cell lines. All four tools except for Numbat performed poorly when applied to cancer cell line alone (**Figure S8**). The inclusion of normal cells improved the performance of these four tools, but not Numbat (**Figure S9**).

In addition, we assessed the performance of the tools in detecting breakpoints. Except for inferCNV, the other four tools were able to output CNV segments at the single-cell level (Figure 3F). Considerable variation in the number of identified segments per cell were observed across these tools. Numbat and CaSpER detected tens to hundreds of segments, while CopyKAT and SCEVAN identified thousands of segments, which clearly exceeded the expected number of chromosomal breakpoints in cancer [27]. Technically it was still challenging to evaluate the breakpoint detection in the levels of single cell. Therefore, we focused our evaluation on the accuracy of the tumor clonal breakpoints in inferCNV, Numbat and SCEVAN (CopyKAT and CaSpER do not implement the function to call clonal breakpoint, see more details in Supplementary Note 2, Figure 3F). CNA calls derived from pseudo bulk scDNA-Seq data were performed using the method adopted in CopyKAT [17] and served as the ground truth. Our results showed that, SCEVAN showed overall the best performance in term of F1 scores and sensitivity, while inferCNV was better at precision (Figure 3G and Figure S10A-B). Both inferCNV and Numbat detected much fewer breakpoints compared with SCEVAN and the ground truth from DNA-Seq (Figure S10C-D). More specifically, the lower F1 score breakpoints for inferCNV and Numbat may be due to reduced resolution in identifying complex CNAs. For instance, SCEVAN detected two breakpoints on chr10, while Numbat and inferCNV merged these two adjacent deletions to one single deletion (Figure 3H and Figure S10E). It could also be attributed to missed breakpoints detection, such as on chr14 (Figure 3I and Figure S10E). Lastly, false positive CNAs calls were also occasionally observed in Numbat (Figure 3J and Figure S10E).

Due to the integration of AB allele analysis, both Numbat and CaSpER are able to detect Loss of Heterozygosity (LOH). The performance of LOH calling for CaSpER was not further evaluated, as it made too many obviously false positive calls both in malignant and nonmalignant cells (Figure S1). Numbat identified a total of 17 LOH events across multiple samples. To assess their accuracy, we then analyzed the actual CNV status of these LOH events using paired scDNA-seq data. In addition, we obtained the merged pseudobulk tumor and normal samples from scDNA-Seq data and performed CNA calling with Sequenza to obtain the allelic copy number states [28]. Our analysis revealed that the majority of LOH events detected by Numbat (14 out of 17, ∼82%) were actually CNV amplifications or deletions (Figure 3K and Figure S11A). For example, in the CRC13 sample, Numbat identified chr8q as an LOH event; however, analysis of CNAs profiles from DNA data revealed that chr8q had actually undergone CNV amplification (Figure 3L). This false positive LOH call by Numbat was due to a significant copy number gain on A allele while the B allele remains unchanged on 8q region, as observed in DNA-Seq analysis (Figure 3M). But Numbat showed high sensitivity in identifying cnLOH. When cross-referenced with DNA data, three regions of chr5q (Figure 3L-M), chr12q (Figure S11A-B), and chr17p (Figure S11C) were identified as cnLOH by Numbat and further validated by AB allele states (B allele copy number is equal to zero for the true positive LOH event) from paired DNA-Seq. Noticeably, the LOH event on chr17p resulted in the loss of heterozygosity of the tumor suppressor *TP53* gene, thus considered to be crucial event during cancer evolution[29, 30]. To further confirm the high sensitivity performance of Numbat on cnLOH calling, we also evaluated one gastric tumor sample (GX109) with four cnLOH events annotated from a published study[31]. All four previously defined cnLOH events were correctly detected by Numbat, and in the meantime no false positive calls were made (Figure S11D). However, considering that false positive LOH calls by Numbat are mostly due to non-neutral copy number status, it is advisable to also reaffirm the neutral copy number state of these regions by independent approaches.

Taken together, the CNAs profiles inferred from scRNA-Seq are overall reasonable, with no single tool clearly outperforming others in terms of CNA inference accuracy. In term of specific tasks, SCEVAN performed the best in breakpoint detection, while Numbat achieved high sensitivity in detecting cnLOH.

### Evaluation of subclonal structure inference accuracy

Human cancer exhibits extensive intra-tumoral heterogeneity, constantly evolving through emerging mutations and CNAs in different subclones, which impact their phenotypes and confer fitness advantages[32]. Both subclones and the tumor evolutionary history can be inferred from single-cell copy number profiles[33]. To evaluate the accuracy of subclonal structure inference by these tools, we selected all cases with more than 10 tumor cells (glioma and 7 CRC samples) (**Figure 1A, TableS1**).

In the case of glioma, previous study has already defined the subclonal structures and only Numbat correctly assigned the smaller C1 subclone (correctly assigning 14 out of 15 cells) with a lower CNAs burden as tumor cells (**Figure 4A-B**). Therefore, we tested the glioma dataset’s subclonal structure inference only using Numbat. We found that the subclone structure inferred by Numbat was largely consistent with scDNA-Seq-based results (**Figure 4C**).

**Figure 4:**
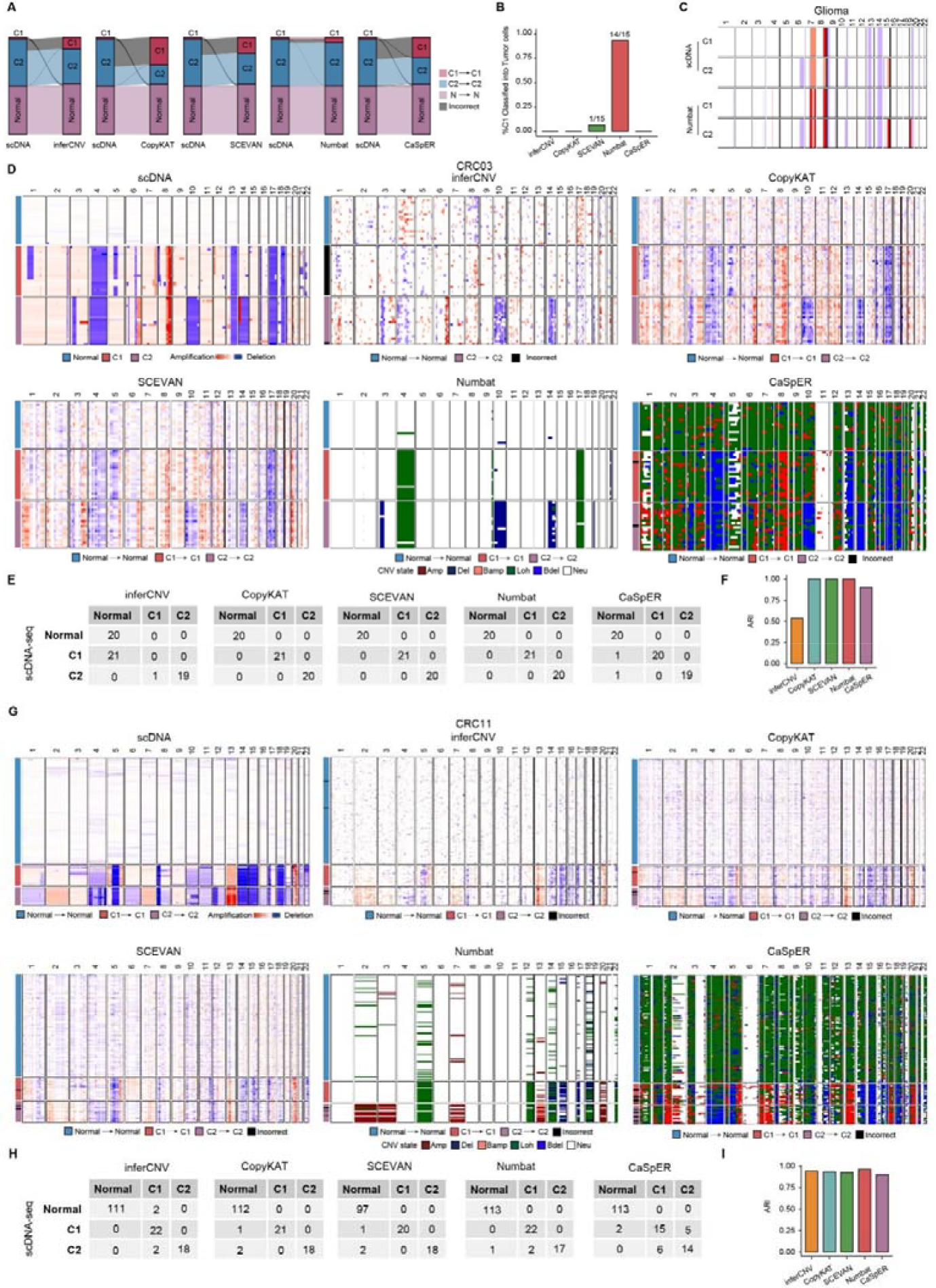
Benchmark of tumor clonal evolution inference. A: Sankey diagrams illustrate the clonal structures inferred from scDNA and scRNA of gliomas. B: A bar plot indicates the percentage of correct classification of the C1 subclone as tumor cells. C: Heatmaps compare the CNAs profiles of two subclones in glioma between scDNA-Seq and those inferred by Numbat. D & G: Heatmaps display the CNAs profiles for CRC03 (D) and CRC11 (G) subclones, obtained from scDNA-seq and scRNA-Seq. E &H: Tables list the cell numbers associated with subclone labels from scDNA and the predicted subclone labels from scRNA for CRC03 (E) and CRC11 (H). F & I: Bar plots present the Adjusted Rand Index (ARI) for subclones in CRC03 (F) and CRC11 (I). Additional information is available in Figure S12-13.

For CRC cases, we studied the cases with at least 10 tumor cells (Figure 2A). First, we obtained the optimal k by average silhouette scores. Interesting, the estimated optimal number of clones for CRC samples were two. If the number of predicted subclones are larger than two by different methods, clusters were combined based on similarity of their CNV profiles (Figure S12A-B). Upon manual inspection, two samples (CRC03 and CRC11) showed two sizeable subclones. In the case of CRC03, a similar tumor calling error occurred only with inferCNV (**Figure 4D**). InferCNV incorrectly assigned the tumor C1 subclone as normal cells, thus failing to detect the two subclones correctly. Adding TME cells to inferCNV corrected the misclassification and improved subclonal structure inference (**Figure S12 C-E**). Most of the other four tools performed well in assigning subclones (**Figure 4 E-F**), although the CNAs profiles for subclones from scDNA-Seq were not fully recapitulated. In the case of CRC11, where all tools correctly classified tumor cells, all tools achieved favorable performance (ARI >0.8 for all tools, **Figure 4G-I**). Interestingly in one extreme case, a subclone defined in CRC12 consisted of only one single cell, and inferCNV and SCEVAN correctly detected such single cell subclone (Figure S13A-B).

To conclude, given the premise of correct tumor cell classification, all five tools achieve desirable performance in delineating the subclonal structures (Figure S13C).

### Aneuploidy inference in normal cells

Accumulating evidence suggest that CNAs can also be found in non-malignant cells and are postulated to exert a crucial impact on cellular fitness[21, 25]. Therefore, we also evaluated the performance of these tools in detecting CNAs in non-malignant cells. Unlike tumor cells with pervasive CNAs across the whole genome, most CNAs observed in non-malignant cells are single whole chromosome gain or loss. We selected four cell types (fibroblast, T, B and endothelial cells) with higher frequent autosomal aneuploidy events[22]. However, the performance of all tools in identifying single-chromosome aneuploidy was generally poor (**Figure S14**). Several factors contributed to such lower detection accuracy. Firstly, CNA complexity in malignant is much higher (compare Figure S1 versus Figure S14). In addition, both the UMI count and gene count per cell in non-malignant cells were significantly lower than that of malignant cells (Figure S15). This highlights the importance of developing more sensitive methods for detecting low-burden CNAs in non-malignant cells.

### Computational speed performance

To evaluate the computation speed of various methods, we tracked the CPU time used (**Table S4**). Our findings indicated that CopyKAT and SCEVAN exhibited the best performance in terms of runtime. InferCNV and CaSpER demonstrated moderate computation speeds, whereas Numbat required the most CPU time. Additionally, we observed a slight increase in runtime with larger datasets. However, the runtime remained acceptable for every tool tested. The calculation of B allele frequency from BAM files is normally the most time as well as resource consuming step, but individual samples can be processed separately in parallel. Based on our tests, a laptop computer could comfortably handle datasets with less than 1000 cells for all five tools. However, for the cases with several thousand cells, analysis is recommended to be executed in a server or high performance computing platform, especially for Numbat and inferCNV.

## Discussion

In this study, we benchmarked five methods used to infer CNAs profiles from scRNA-seq data. Overall, we observed that all five tools could distinguish between tumor and normal cells and accurately infer CNAs profiles to a certain extent. Among these methods, Numbat demonstrated the best performance across various evaluation criteria. For case where only the expression matrix was available, CopyKAT is recommended as the preferred method. Additionally, we have included a comprehensive and informative table (**Figure 5**) that summarizes the characteristics of all the benchmarked methods.

**Figure 5:**
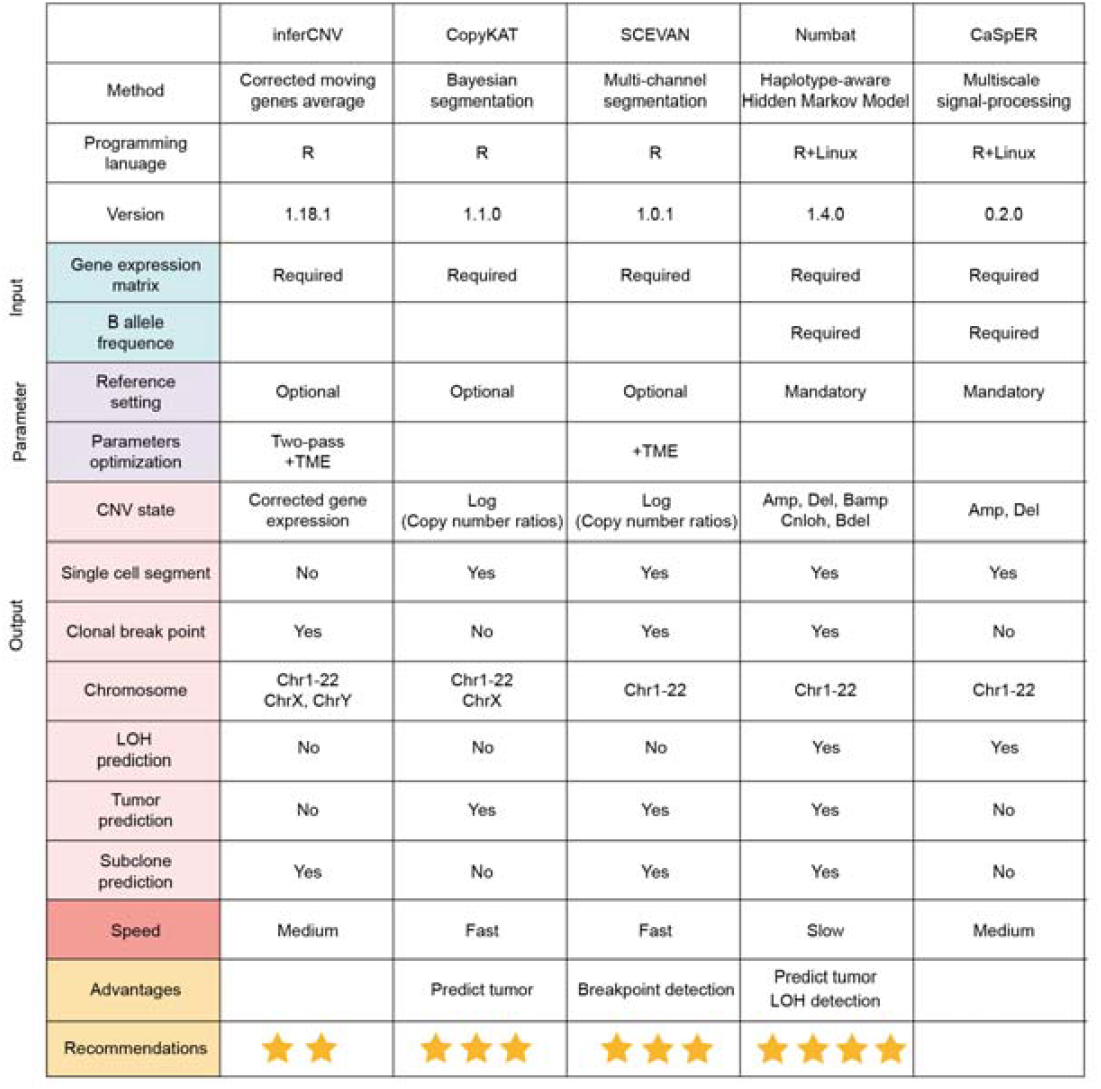
Comparison overview for all methods. We systematically summarized the methodology, programming language used, tools version, input data formats, parameters settings, output data, computational speed and recommendations for each software. Two-pass represents running inferCNV twice and inputs normal cell inferred from the first run as reference cells, see Figure3A for more details; +TME: including tumor microenvironment cells as input.

Methods based on gene expression are more susceptible to the influence of tumor purity. Without a reference setting, inferCNV relies on the input cell population to generate a background. When tumor purity is high, actual CNAs signals may be incorrectly considered as background and removed. Consequently, inferCNV occasionally exhibited incorrect centering of CNAs, leading to mis-classification of normal and tumor cells in our benchmark (**Figure 2D and S4**). On the contrary, SCEVAN tended to incorrectly assign non-malignant cells as malignant cells when tumor purity was low. Interestingly, including TME cells in the analysis improved performance in both cases (**Figure 2E, Figure S3A**,**C**).

Incorporation of B allele information enabled Numbat and CaSpER to identify LOH events, which play crucial roles in cancer progression[28]. Our results showed that Numbat showed high sensitivity but low precision in LOH detection. Most of false positive cnLOH calls were found to be copy number deletion (Figure 3K), which failed to be classified into copy number deletion due to the model’s resolution to distinguish (1,0) and (2,0) genotypes. The notation (2,0) indicates a copy number neutral LOH, where the cell contains two copies of the A allele and no copies of the B allele. This means that the B allele is lost, but the total copy number remains unchanged (2 copies of A). On the other hand, (1,0) represents a copy number deletion-mediated LOH, where the cell contains one copy of the A allele and no copies of the B allele [19]. Although copy number gain normally shouldn’t result in LOH, as both A and B allele DNA remain present. Extreme cases of copy number amplification (such as the example on chr8 in Figure 3L-M) were also mis-classified as LOH by Numbat. This was again likely due to the genotype modeling setup in Numbat. Because the most imbalanced genotype for copy number gain is three copy of A allele and one copy of B allele (3,1), which might be considered less likely compared with cnLOH associated (2,0) genotype. Therefore, users should be careful in interpreting the LOH results from Numbat. It is recommended to confirm the absence of copy number alterations in these LOH regions using independent tools.

Another observation from our comparative analysis was that the complexity and burden of CNAs affected each method equally. We found that tools generally performed better on solid tumors than on hematopoietic cancers or cell lines. This could be due to the fewer copy number alterations and lower chromosomal CNAs heterogeneity in hematopoietic cancers. Additionally, the methods performed poorly in identifying aneuploidy in nonmalignant cells. This underscored the need to develop more sensitive methods for inferring low-burden CNAs from scRNA-seq to address this gap in the field. On the contrary, another systematic factor--the sequencing depth affects the performance of five tools differently. In this case, the dependence of B allele information also came with a cost. Compared with inferCNV or CopyKAT, the performance of Numbat deteriorated significantly if the UMI counts per cell fell below 10k (Figure 2J).

Given no single tool excelled in every task, we recommended users to perform the analysis with at least two tools. When raw data is available, Numbat and SCEVAN (or inferCNV) combination can be considered. CopyKAT and SCEVAN (or inferCNV) combination is recommended for cases with expression matrix alone.

One limitation of our benchmark study was the limited availability of single cell multiomics datasets that simultaneously measured both DNA and RNA in the same cells. Therefore, the number of input cells for CNAs inference in these datasets is relatively small compared to microfluidics-based scRNA-Seq approaches such as 10X Genomics platform. Therefore, we were unable to evaluate how different method perform for large and more complex datasets. In addition, the small number of cells also limits our evaluation of subclonal structure inference. Last but not least, the parameter setting plays a crucial role in the performance, especially for the ones with more complex modeling such as Numbat, which offers a larger number of tunable parameters compared to other tools. We sought to optimize the parameter settings for each tool by combining insights from the literature, recommendations from GitHub documentation, and our own testing across the entire dataset. But we observed that the optimized parameters varied depending on both the sample characteristics and the specific tasks. For instance, a smaller gamma parameter in Numbat yielded better tumor/normal distinction, but compromised their copy number breakpoint detection in some samples. Therefore, we highly recommended the users to try to adjust key parameters and perform few runs to achieve optimal performance from these tools.

In summary, this study provides a much-needed independent comparison of CNAs inference tools. Our results will help researchers gain a deeper understanding of the performance and limitations of current methodologies, enabling them to choose and use these tools appropriately. Finally, we believe that many of the lessons learned in this benchmark study may also apply to CNAs inference in spatial transcriptomics.

## Key Points

- Single cell multi-omics datasets (profiled DNA and RNA in the same cell) were used as benchmark datasets.
- We assessed five state-of-the-art scRNA-Seq based SCNA inference methods and found that Numbat performed the best and showed high sensitivity in cnLOH detection.
- CopyKAT demonstrated superior performance when using expression matrices alone as input and showed robustness in lower sequencing depth.
- SCEVAN achieved the best results in clonal breakpoint detection.
- We provided recommendations on how to set parameters to optimize different tools’ performance.

## Methods

### Benchmark datasets collection

The CRC dataset was sourced from the study by Zhou et al.[22]. Both raw FASTQ files and processed gene expression count matrices for scRNA-seq were downloaded from the Genome Sequence Archive of the National Genomics Data Center (NGDC) with the accession number HRA000201. The glioma, NPC43 and HUVEC cell line data were obtained from the study by Yu et al[24] with gene expression count matrices for scRNA-seq available from the NCBI Gene Expression Omnibus under accession number GSE185269. The Neuro dataset was sourced from the study by Cui et al[26]. Both raw FASTQ files and processed gene expression count matrices for scRNA-seq were downloaded from the Genome Sequence Archive of the National Genomics Data Center (NGDC) with the accession number PRJCA002946. The GX109 dataset was sourced from the study by Huang et al[31]. Raw FASTQ data were aligned to the genome to generate BAM files, following the processing methods described in the original articles by Zhou et al, Yu et al and Cui.[22, 24, 26]. Additionally, the processed count matrix for two acute lymphoblastic leukemia samples was obtained from the study by Zachariadis et al with accession number GSE144296[23]. Chromosomal copy number variations in scDNA-seq data and cell type annotations for all these datasets was acquired directly from the authors.

### Workflows for inferCNV[16], CopyKAT[17], SCEVAN[18], Numbat[19] and CaSpER[20] InferCNV

It utilizes a corrected moving average technique to compute expression levels of genes that are adjacent in the chromosomes. This approach helps mitigate gene-specific expression variability and generates chromosomal copy number profiles. The genes are arranged based on their absolute genomic positions, initially organized by chromosome and subsequently by the genomic start position within each chromosome. Leveraging the moving average expression data, InferCNV generates CNV profiles for samples, representing inferred copy number variations across diverse genomic regions. To further refine the CNV profiles of tumor cells, CNAs profiles were adjusted by subtracting the corresponding values from either normal cells or the average of all cells. In this project, we followed the guidelines on the GitHub repository of inferCNV (https://github.com/broadinstitute/infercnv). inferCNV version 1.18.1 was executed with default parameters, except for the ‘tumor_subcluster_partition_method’ parameter, which was set to ‘random_trees’.

### CopyKAT

It combined Bayesian approach with hierarchical clustering to infer copy number profiles of individual cells and delineate clonal substructure from a matrix of unique molecular identifier (UMI) counts. Similar to the inferCNV method, CopyKAT begins by arranging genes based on their genomic coordinates. It then employs integrative hierarchical clustering and Gaussian mixture modeling (GMM) to identify potentially normal cells with high confidence, thereby inferring the copy number (CN) baseline. The subsequent step involves detecting chromosome breakpoints where CNA profiles exhibit changes. To accomplish this, CopyKAT integrates a Poisson-gamma model with Markov chain Monte Carlo (MCMC) iterations to compute posterior means per genomic window and employs Kolmogorov–Smirnov (KS) tests to connect adjacent windows. Ultimately, the final copy number values for each window are determined as the posterior averages of all genes spanning the adjacent chromosome breakpoints in each cell. In this project, we followed the guidelines provided on the GitHub repository for CopyKAT (v 1.1.0) (https://github.com/navinlabcode/copykat). CopyKAT was used with default parameters.

### SCEVAN

It is capable of automatically distinguishing between tumor and normal cells and inferring copy number breakpoints using an optimization-based joint segmentation algorithm. The analysis begins with a raw count matrix for genes, which is first log-transformed and filtered to remove genes detected in less than 10% of cells. Subsequently, a subset of high-confidence normal cells is then identified using gene signatures through a selection of those from the tumor microenvironment, stromal cells, and immune cells. The median expression of these normal cells serves as the baseline for copy number estimation. Next, a relative gene expression matrix is then derived by subtracting this baseline, followed by edge-preserving nonlinear diffusion filtering to achieve smoothed expression profiles. These smoothed expression profiles are then subjected to multichannel segmentation based on the Mumford and Shah energy principle, resulting in to a copy number matrix. In this project, we followed the guidelines provided in the SCEVAN (v 1.0.1) GitHub repository (https://github.com/AntonioDeFalco/SCEVAN). SCEVAN was executed using the default parameters, with the exception of setting ‘plotTree’ to ‘FALSE’ in order to reduce the running time.

### Numbat

It integrates expression, allele and haplotype information derived from population-based phasing to comprehensively characterize copy number variations profiles. The workflow of Numbat involves two key steps. First, it extracts allele counts at the cell level from BAM format files. Subsequently, it employs a haplotype-aware Hidden Markov Model (HMM) to infer CNVs within cell population pseudobulk profiles. The assignment of posterior probabilities to cells is used to determine whether a particular cell exhibits specific copy number alterations. In this project, we followed the guidelines provided on the GitHub repository of Numbat: https://kharchenkolab.github.io/numbat/articles/numbat.html. We obtained allele data using the default parameters of the pileup_and_phase.R script. We next used the run_numbat function from the Numbat package (version 1.3.2.1) to obtain CNA profiles with the default parameters. For Non-UMI protocols data, we set the expectation gamma parameter to 5, based on the author’s recommendation. However, due to the severe imbalance in the number of tumor cells and normal cells in CRC22 and CRC15, Numbat cannot run successfully with the default parameters. To address this, we made specific adjustments. For the CRC22 dataset, we set the min_LLR parameter to 1, and the max_entropy to 0.6. For the CRC15 dataset, we set the min_cells parameter to 1 in order to obtain more filtered CNAs segments.

### CaSpER

It utilizes a multiscale signal processing framework that integrates two streams of information: gene expression and B-allele frequencies (BAF), derived from either bulk or single-cell RNA sequencing data. Initially, CaSpER applies recursive iterative median filtering to smooth both the BAF and expression signals. At each scale, a hidden Markov model is employed to analyze the smoothed expression signal and categorize it into five CNAs states: (1) homozygous deletion, (2) heterozygous deletion, (3) neutral, (4) one-copy amplification, and (5) multi-copy amplification. For segments labeled as states 2 and 4, CaSpER incorporates information from the BAF signal. If a shift in the BAF signal accompanies the segment, states 2 and 4 are adjusted to states 1 and 5, respectively. BAF shifts are determined by thresholding the smoothed signal, with thresholds calculated using a Gaussian mixture model fit on pooled samples across a segment. Consequently, CaSpER provides discrete CNA classifications: amplification, neutral, or deletion, as a result of these thresholds and corrections. In this project, we followed the guidelines on the GitHub repository of CaSpER: https://github.com/akdess/CaSpER. CaSpER (v 0.2.0) was executed using default parameters. Both required parameters (“cnv.scale” and “loh.scale”) were set to 3 based on example codes.

### Generation of synthetic datasets by downsampling of full glioma dataset spanning full range of tumor vs normal ratios

We used R (version 4.3.2) to downsample the glioma data based on the cell number ratio between tumor and normal cells of 1:100, 1:20, 1:4, 1:2, 2:1, 4:1, 20:1, and 100:1. Each of these tumor/normal ratios was resampled 50 times. The ratio of 1:1 represents the original full data. For each synthetic dataset, inferCNV, CopyKAT, SCEVAN, Numbat, and CaSpER software were employed to classify tumor and normal cells, and F1 scores were computed. The mean and standard error of F1 scores of 50 resamplings at each condition are reported.

### Simulation of down-sampled data with fixed sequencing depths

We used samtools (version 1.8) [34] to randomly subsample reads from the BAM files with the -s option to specify the random seed and proportion with samtools view -s random_seed ratio. Each sample was randomly subsampled three times, with the random seeds set to 555, 666, and 777, and subsequently calculated the UMI count for each cell, following the processing methods described in the original study by Zhou et al[22]. This approach ensured that the median UMI per cell for each sample was approximately 10K, 3K, and 1K, simulating different levels of sequencing depth. The simulated datasets were processed similarly as the original dataset.

### Reference setting for Numbat and CaSpER

Numbat and CaSpER tools must specify reference expression. For Numbat, the parameter ‘lambdas_ref’ was used to regress out any cell-type-specific expression differences unrelated to CNVs. The reference expression for CRC and fibroblasts data was sourced from non-epithelial cells of corresponding patients. For glioma data, it was originated from astrocyte, oligodendrocyte, GABAergic, and glutamatergic cells. The reference expression for ALL was derived from TME cells. The reference expression for NPC43 cell line data was derived from the HUVEC cell line. For CaSpER, except for CRC data, other data used reference cells consistent with Numbat. Non-epithelial cells were not used in CRC data because we aimed to distinguish potential normal epithelial cells from tumor epithelial cells as reference cells to further improve the accuracy of CaSpER. This approach was more accurate than using non-epithelial cells (such as immune cells, fibroblasts, endothelial cells), which have cell type-specific transcription profiles that compromise the accuracy of CNAs detection. Therefore, we first divided epithelial cells into two categories through the Louvain algorithm. Generally, the transcriptional activity of normal epithelial cells is lower than that of tumor cells, resulting in lower UMIs. Hence, we considered the subset of cells with lower UMIs as the reference. For fibroblasts, we used the mean values across cells as reference for CaSpER. We’ve also included a detailed guide for reference selection in Supplementary Note 3.

### Classification of tumor and normal cells

For CopyKAT, SCEVAN, and Numbat, we utilized the software’s inherent classification to distinguish between tumor and normal cells. Since inferCNV and CaSpER do not directly classify tumor cells, we conducted hierarchical clustering on individual cell CNAs profiles, aiming for two clusters. We then determined the cluster with the higher standard deviation in mean copy number across cells as tumor cells, with the other cluster being normal cells.

### Evaluation CNAs profile accuracy

We evaluated the accuracy of CNA profiles by calculating the Pearson correlation between scDNA-seq and bulk DNA-seq data. The heatmap of CNAs represents the median CAN values from individual cells. As Numbat and CaSpER provide categorical outputs, we assigned numerical equivalents: “deletion” as -1, “neutral” as 0, and “amplification” as 1.

### Evaluation of clonal breakpoint detection accuracy

Pseudo-bulk CNV breakpoints were estimated from scDNA-seq data as the ground truth, following an approach similar to that described in the CopyKAT. Briefly, pseudo-bulk CNV breakpoints were obtained by the circular binary segmentation (CBS) method using DNAcopy (version 1.68.0) package[35] to segment genomic bins, followed by MergeLevels [36] to merge adjacent segments with nonsignificant differences in segment ratios. A clonal breakpoint detected in scRNA-seq data was considered a True Positive (TP) if it was located within 200 kb from the true breakpoints identified in scDNA-seq. A False Negative (FN) was recorded when no scRNA-Seq breakpoint was detected within 200 kb of scDNA-seq defined breakpoint. A False Positive (FP) was defined as a breakpoint identified in scRNA-seq but not supported by within 200kb scDNA-seq defined breakpoints. We evaluated the F1 score, precision and sensitivity for inferCNV, SCEVAN and Numbat, as only these three tools could output clonal CNA breakpoints.

### Evaluation of LOH detection accuracy

Following the processing described in the original article, the raw FASTQ data is aligned with the genome to generate the BAM file[22]. According to the barcode information corresponding to Tumor cells and normal cells (normal epithelial + TME) cells, the BAM files of individual Tumor cells and normal cells were separated from each DNA library and merged into pseudo-bulk tumor and pseudo-bulk normal samples separately. Then AB allele calling was performed by Sequenza to verify the accuracy of LOH event [37].

### Construction Clonal evolution inference

Subclonal structures defined by paired scDNA-Seq served as ground truth. Considering several tools such as inferCNV, CopyKAT and Numbat, subclones are obtained based on hierarchical clustering of CNAs profiles. In order to establish a uniform evaluation standard, we also performed hierarchical clustering by Euclidean distance with Ward.D2 linkage on CNA profiles. We obtained the optimal number of clusters (k) using the average silhouette scores.

Three (inferCNV, Numbat and SCEVAN) out of five tools had built-in function to infer subclonal structure. However, we frequently encountered bugs in execution when using SCEVAN for subclonal structure calling where the number of subclones in certain samples was low. Consequently, for the purposes of this study, we performed subclone identification using scDNA-Seq based subclonal structure calling pipeline for CopyKAT, SCEVAN and CaSpER based on their cell-by-segmentation output matrixes. We obtained the subclonal structure output from inferCNV and Numbat directly. When more than two subclones were predicted in these two methods, clusters were combined based on the similarity of their CNV profiles (Figure S12AB).

Lastly, to match scRNA-seq derived subclones with those from scDNA-seq, we calculated the median values for each subclone and computed the Pearson correlation. Subclones were assigned based on the highest correlation.

### Aneuploidy cells identification in TME cells

Fibroblasts, T cells, B cells and endothelial cells were selected for analysis. All aneuploidy and diploidy cell annotations were obtained from the Zhou et al[22]. We calculated the ability of F1 scores, Sensitivity and Specificity for identifying autosomal aneuploid cells (aneuploidy in X or Y chromosomes were not included) to evaluate five tools.

### Genome-level heatmap

Due to inconsistencies in the start and end coordinates of CNAs segments on chromosomes across different software outputs, we employed the following formula to uniformly compute the average signal by splitting the genome into 1MB windows.

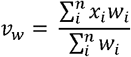

*v*_*W*_ represents the weighted mean of all signal regions, with the weights determined by the width of their intersections in the 1MB window. *w*_*l*_ denotes the interaction width of CNAs segments within the 1MB window, and *x*_*l*_ represents the corresponding CNAs signal value.

### Computational speed performance

We evaluated both time and memory usage on a High-Performance Computing platform. Each method was executed using a single-core setting. Runtime was captured for each method using the Sys.time() function in R (v.4.1.2).

### Data visualization

Heatmaps were visualized by R package ComplexHeatmap (v2.10.0)[38]. All bar plots and box plots were visualized using the R packages ggpubr (v 0.4.0)[39].

### Statistics

All statistical analyses and presentations were performed using R (v4.1.2). The methods of statistical test were indicated in the figure legends, and adjustments for multiple comparisons were made using the Benjamini & Hochberg method when necessary.

## Declarations

## Ethics approval and consent to participate

Not applicable.

## Consent for publication

Not applicable.

## Availability of data and materials

All the scRNA-seq datasets used in this study are publicly available datasets, and their accession numbers could be found in Table S1. The code and scripts used for the benchmark analysis and figure plotting has been uploaded to GitHub: https://github.com/bioliyezhang/Benchmark_scRNASeq_by_scMultiomics.

## Authors’ contributions

L.Z., M.S., Y.Z. and S.M. wrote the paper. L.Z. designed the study with M.S. and S.M. M.S. and S.M. wrote code, prepared the data, analysis and figures. G. W and Y.W. performed analysis. L.Z. and X.H. supervised the work. B.X. and T.G. modified the details of the figures. All authors reviewed and approved the final paper.

## Acknowledgements

We thank Prof. Fuchou Tang from Peking University, Prof. Shuhui Bian from Nanjing Medical University, Prof. Angela Ruohao Wu and Dr. Lei Yu from Hong Kong University for providing data. We also thank ShanghaiTech University’s high-performance computing platform and Zhejiang lab for support.

## Reference

1. Wen L, Li G, Huang T et al. Single-cell technologies: From research to application, Innovation (Camb) 2022;3:100342.

2. Jovic D, Liang X, Zeng H et al. Single-cell RNA sequencing technologies and applications: A brief overview, Clin Transl Med 2022;12:e694.

3. Andrews TS, Kiselev VY, McCarthy D et al. Tutorial: guidelines for the computational analysis of single-cell RNA sequencing data, Nat Protoc 2021;16:1–9.

4. Xue R, Zhang Q, Cao Q et al. Liver tumour immune microenvironment subtypes and neutrophil heterogeneity, Nature 2022;612:141–147.

5. Ren X, Zhang L, Zhang Y et al. Insights Gained from Single-Cell Analysis of Immune Cells in the Tumor Microenvironment, Annu Rev Immunol 2021;39:583–609.

6. Sun Y, Wu L, Zhong Y et al. Single-cell landscape of the ecosystem in early-relapse hepatocellular carcinoma, Cell 2021;184:404–421 e416.

7. Peng J, Sun BF, Chen CY et al. Single-cell RNA-seq highlights intra-tumoral heterogeneity and malignant progression in pancreatic ductal adenocarcinoma, Cell Res 2019;29:725–738.

8. Puram SV, Tirosh I, Parikh AS et al. Single-Cell Transcriptomic Analysis of Primary and Metastatic Tumor Ecosystems in Head and Neck Cancer, Cell 2017;171:1611–1624 e1624.

9. Chung W, Eum HH, Lee HO et al. Single-cell RNA-seq enables comprehensive tumour and immune cell profiling in primary breast cancer, Nat Commun 2017;8:15081.

10. Bailey C, Black JRM, Reading JL et al. Tracking Cancer Evolution through the Disease Course, Cancer Discov 2021;11:916–932.

11. Knouse KA, Wu J, Amon A. Assessment of megabase-scale somatic copy number variation using single-cell sequencing, Genome Res 2016;26:376–384.

12. Baslan T, Kendall J, Ward B et al. Optimizing sparse sequencing of single cells for highly multiplex copy number profiling, Genome Res 2015;25:714–724.

13. Cai X, Evrony GD, Lehmann HS et al. Single-cell, genome-wide sequencing identifies clonal somatic copy-number variation in the human brain, Cell Rep 2014;8:1280–1289.

14. Ruohan W, Yuwei Z, Mengbo W et al. Resolving single-cell copy number profiling for large datasets, Brief Bioinform 2022;23.

15. Garvin T, Aboukhalil R, Kendall J et al. Interactive analysis and assessment of single-cell copy-number variations, Nat Methods 2015;12:1058–1060.

16. Tirosh I, Izar B, Prakadan SM et al. Dissecting the multicellular ecosystem of metastatic melanoma by single-cell RNA-seq, Science 2016;352:189–196.

17. Gao R, Bai S, Henderson YC et al. Delineating copy number and clonal substructure in human tumors from single-cell transcriptomes, Nat Biotechnol 2021;39:599–608.

18. De Falco A, Caruso F, Su XD et al. A variational algorithm to detect the clonal copy number substructure of tumors from scRNA-seq data, Nat Commun 2023;14:1074.

19. Gao T, Soldatov R, Sarkar H et al. Haplotype-aware analysis of somatic copy number variations from single-cell transcriptomes, Nat Biotechnol 2023;41:417–426.

20. Serin Harmanci A, Harmanci AO, Zhou X. CaSpER identifies and visualizes CNV events by integrative analysis of single-cell or bulk RNA-sequencing data, Nat Commun 2020;11:89.

21. Erickson A, He M, Berglund E et al. Spatially resolved clonal copy number alterations in benign and malignant tissue, Nature 2022;608:360–367.

22. Zhou Y, Bian S, Zhou X et al. Single-Cell Multiomics Sequencing Reveals Prevalent Genomic Alterations in Tumor Stromal Cells of Human Colorectal Cancer, Cancer Cell 2020;38:818–828 e815.

23. Zachariadis V, Cheng H, Andrews N et al. A Highly Scalable Method for Joint Whole-Genome Sequencing and Gene-Expression Profiling of Single Cells, Mol Cell 2020;80:541–553 e545.

24. Yu L, Wang X, Mu Q et al. scONE-seq: A single-cell multi-omics method enables simultaneous dissection of phenotype and genotype heterogeneity from frozen tumors, Sci Adv 2023;9:eabp8901.

25. Iourov IY, Vorsanova SG, Yurov YB. Somatic genome variations in health and disease, Curr Genomics 2010;11:387–396.

26. Cui Y, Li C, Jiang Z et al. Single-cell transcriptome and genome analyses of pituitary neuroendocrine tumors, Neuro Oncol 2021;23:1859–1871.

27. Cornish AJ, Gruber AJ, Kinnersley B et al. The genomic landscape of 2,023 colorectal cancers, Nature 2024;633:127–136.

28. Ciani Y, Fedrizzi T, Prandi D et al. Allele-specific genomic data elucidate the role of somatic gain and copy-number neutral loss of heterozygosity in cancer, Cell Syst 2022;13:183–193 e187.

29. Baslan T, Morris JPt, Zhao Z et al. Ordered and deterministic cancer genome evolution after p53 loss, Nature 2022;608:795–802.

30. Murai K, Dentro S, Ong SH et al. p53 mutation in normal esophagus promotes multiple stages of carcinogenesis but is constrained by clonal competition, Nat Commun 2022;13:6206.

31. Huang R, Huang X, Tong Y et al. Robust analysis of allele-specific copy number alterations from scRNA-seq data with XClone, Nat Commun 2024;15:6684.

32. Dentro SC, Leshchiner I, Haase K et al. Characterizing genetic intra-tumor heterogeneity across 2,658 human cancer genomes, Cell 2021;184:2239–2254 e2239.

33. Wang F, Wang Q, Mohanty V et al. MEDALT: single-cell copy number lineage tracing enabling gene discovery, Genome Biol 2021;22:70.

34. Danecek P, Bonfield JK, Liddle J et al. Twelve years of SAMtools and BCFtools, Gigascience 2021;10.

35. Seshan VE OA. DNAcopy: DNA Copy Number Data Analysis. R package version 1.81.0. https://bioconductor.org/packages/DNAcopy.

36. Willenbrock H, Fridlyand J. A comparison study: applying segmentation to array CGH data for downstream analyses, Bioinformatics 2005;21:4084–4091.

37. Favero F, Joshi T, Marquard AM et al. Sequenza: allele-specific copy number and mutation profiles from tumor sequencing data, Ann Oncol 2015;26:64–70.

38. Gu Z, Eils R, Schlesner M. Complex heatmaps reveal patterns and correlations in multidimensional genomic data, Bioinformatics 2016;32:2847–2849.

39. Kassambara A. ggpubr: ‘ggplot2’ Based Publication Ready Plots. https://CRAN.R-project.org/package=ggpubr.

