## Supplemental Files for "Benchmarking copy number aberrations inference tools using single-cell multi-omics datasets"

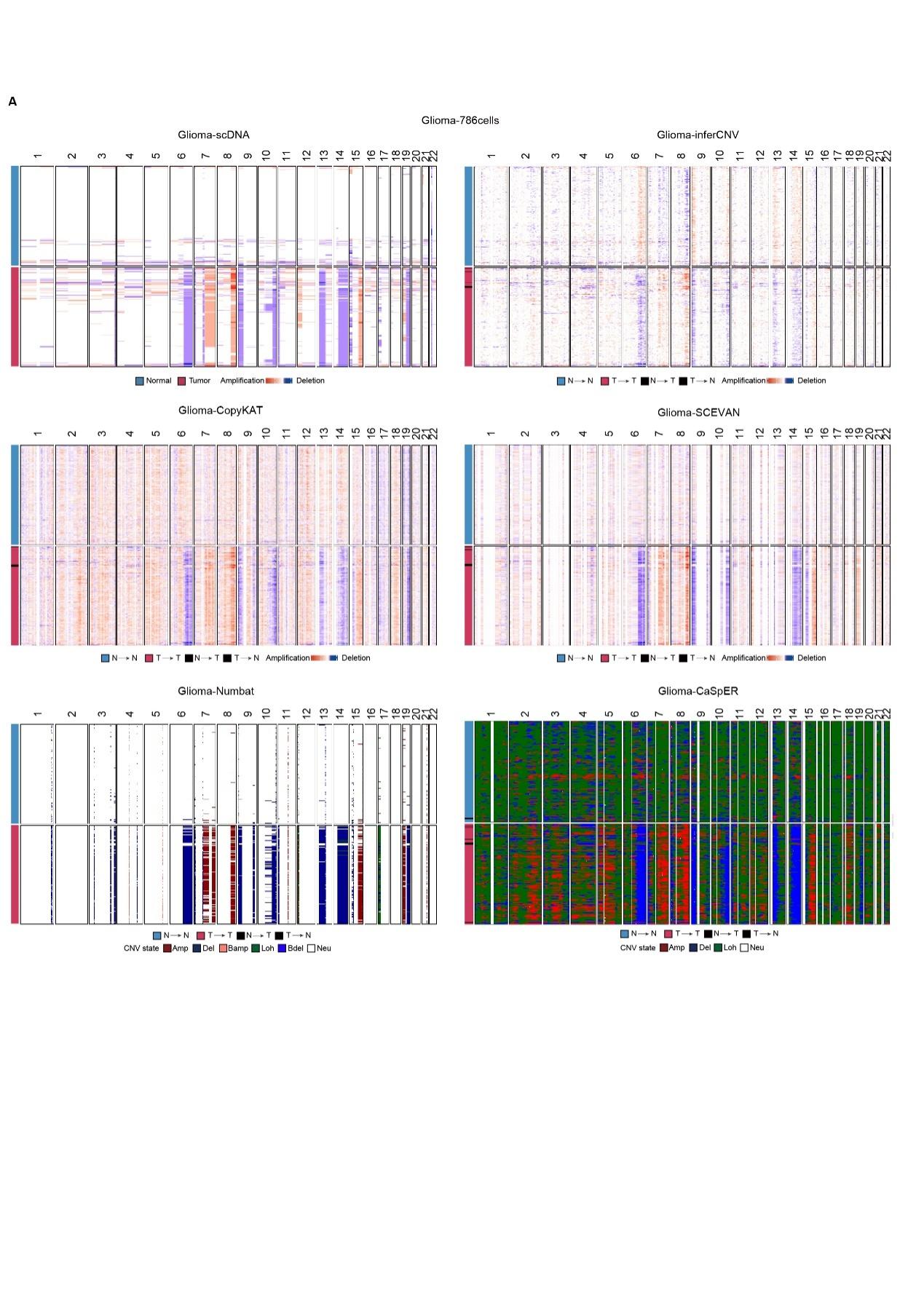

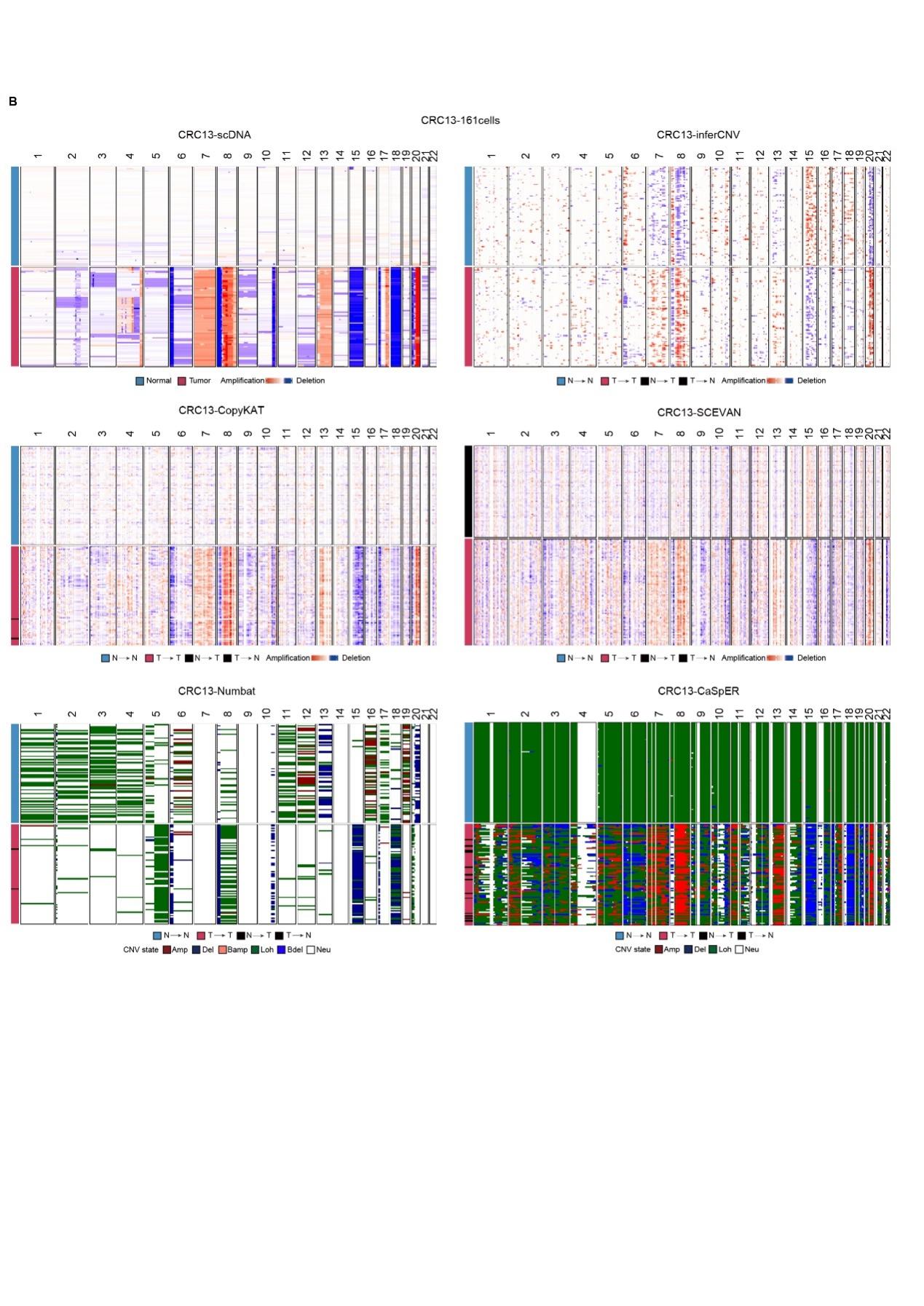

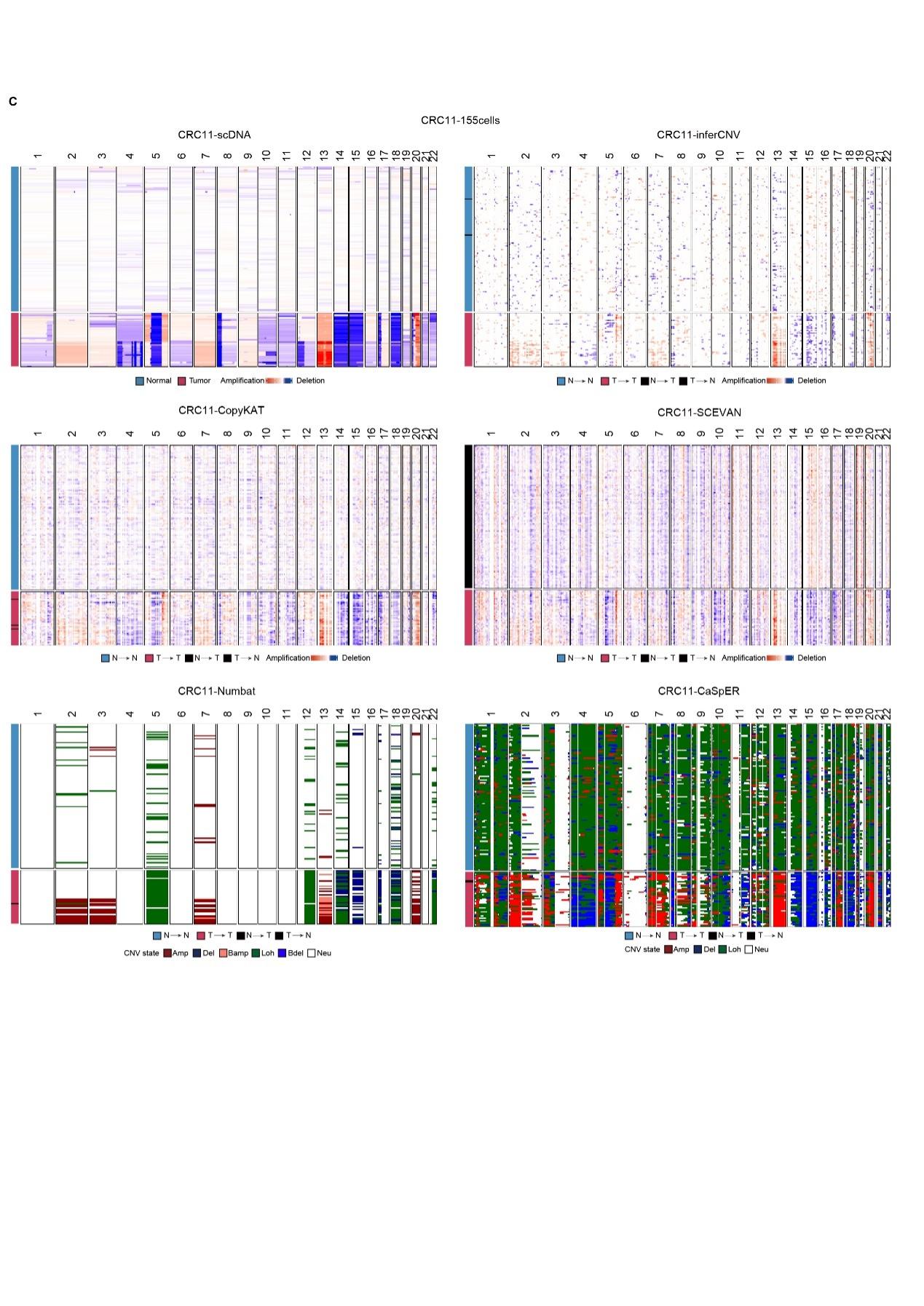

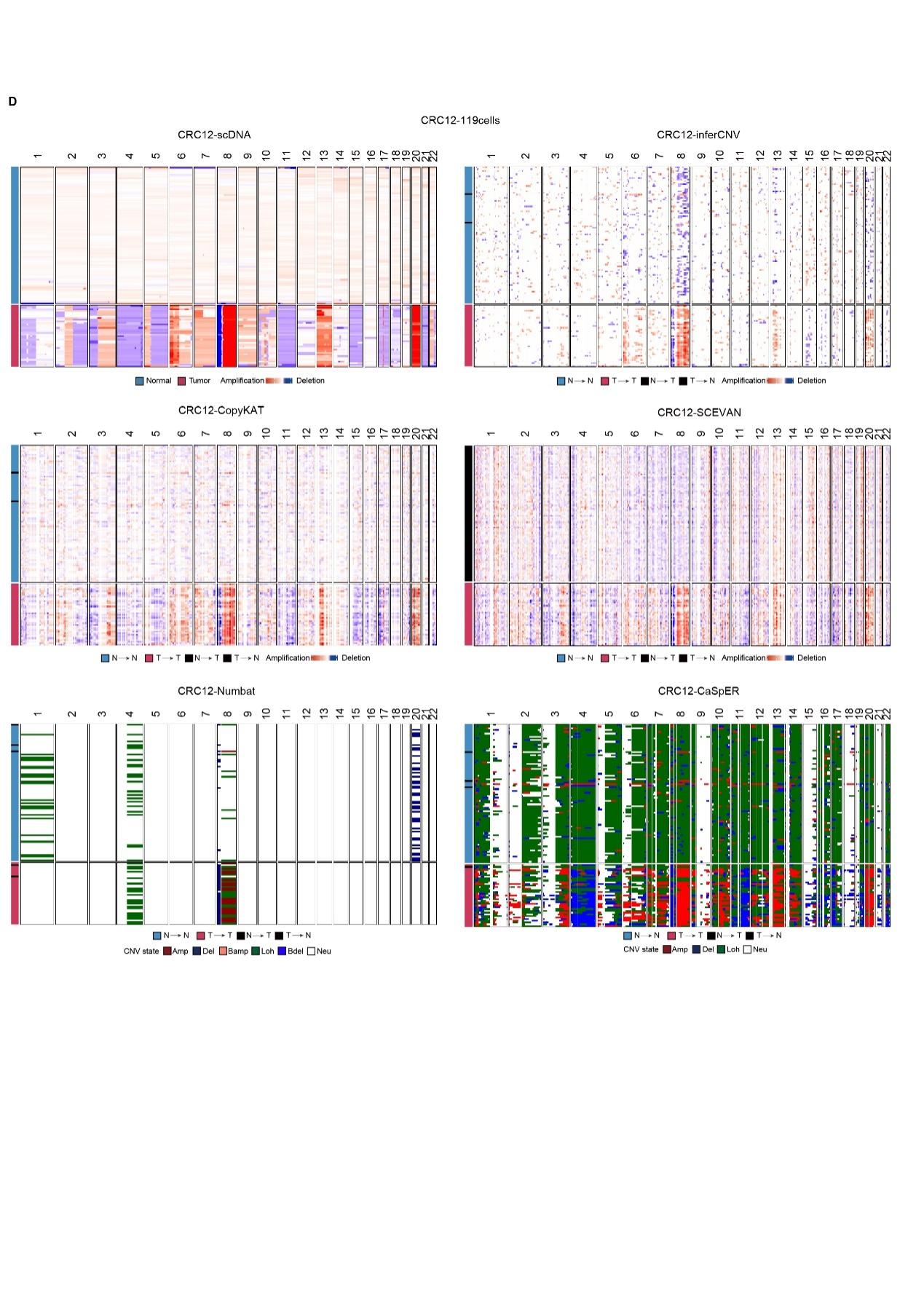

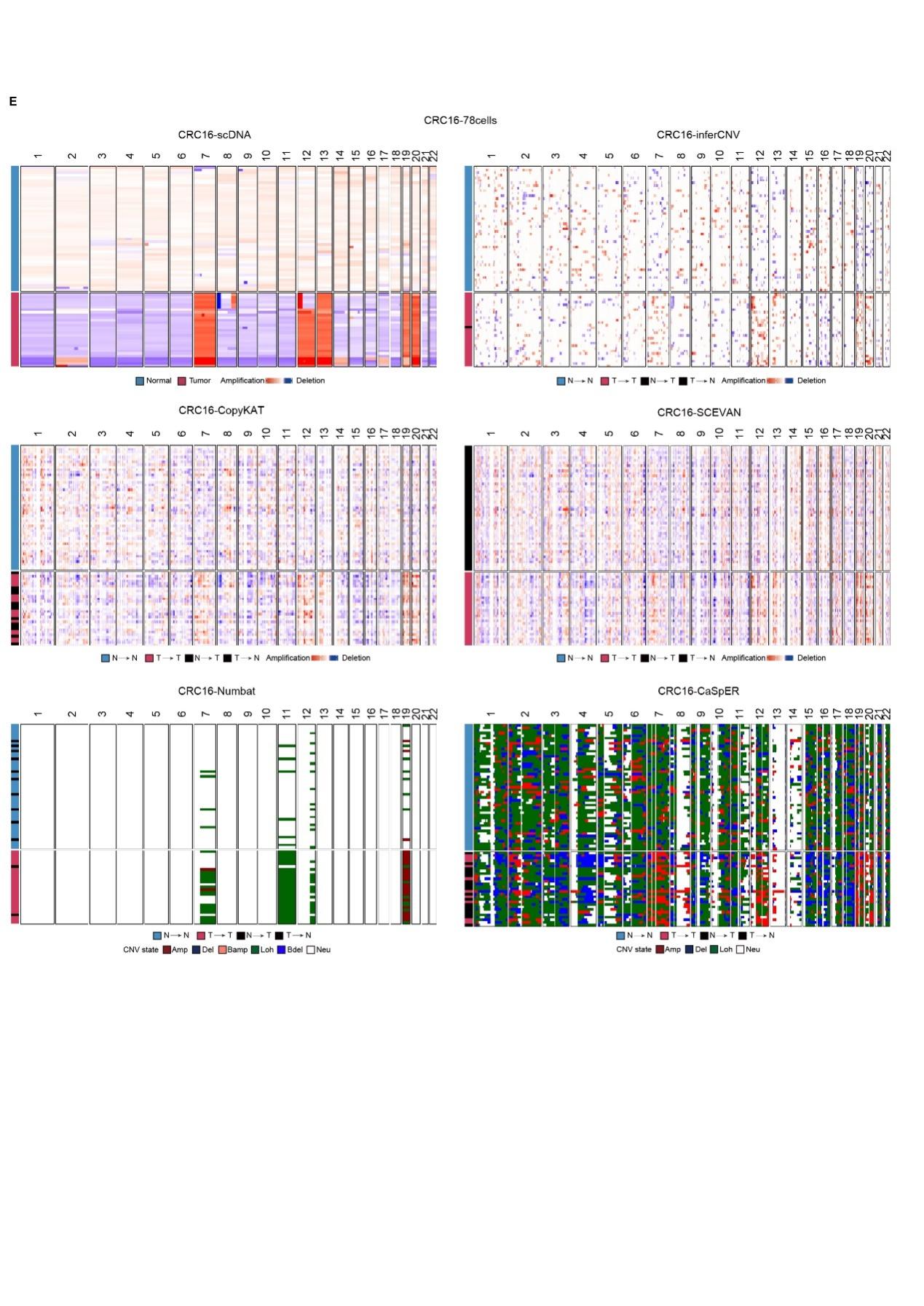

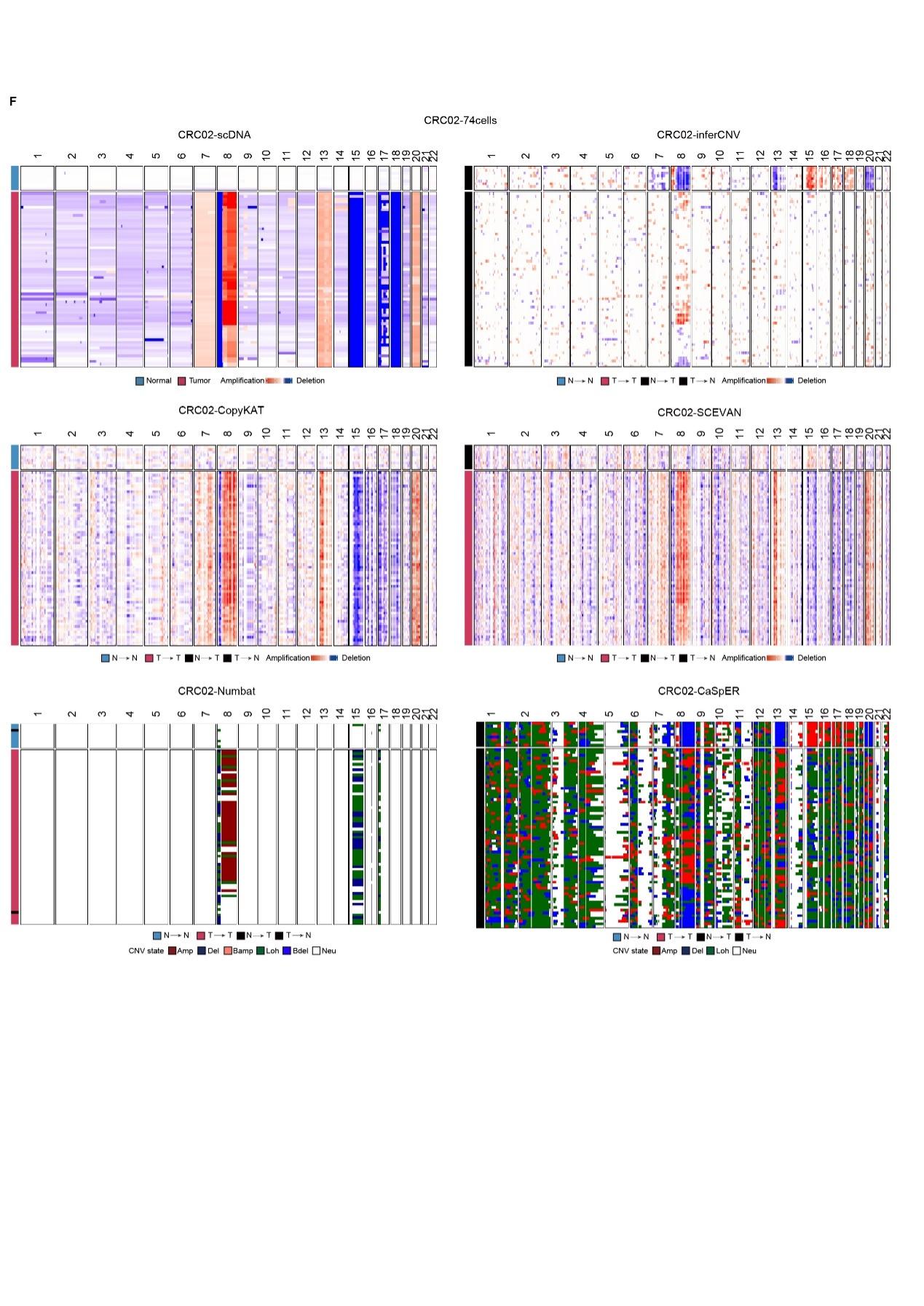

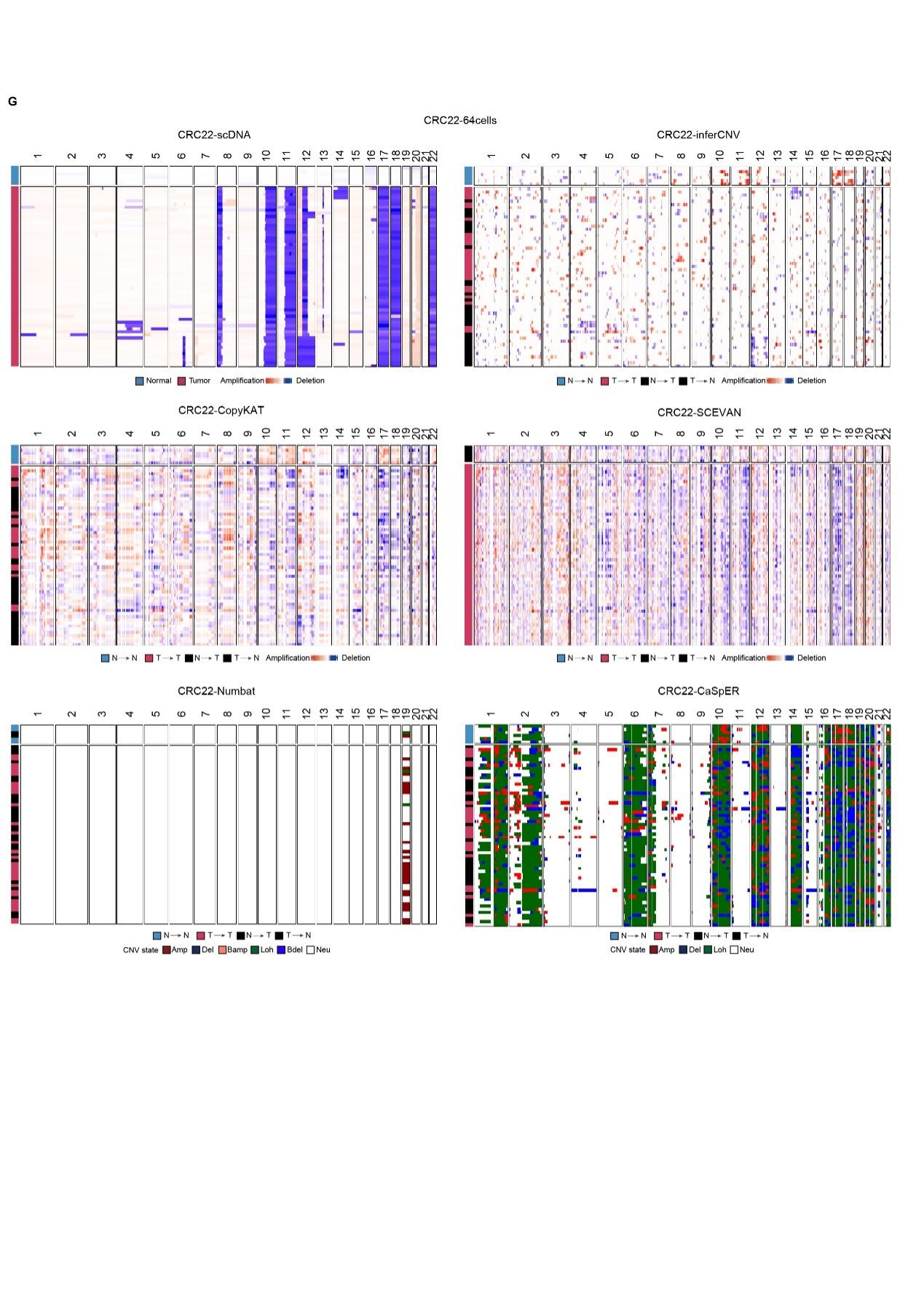

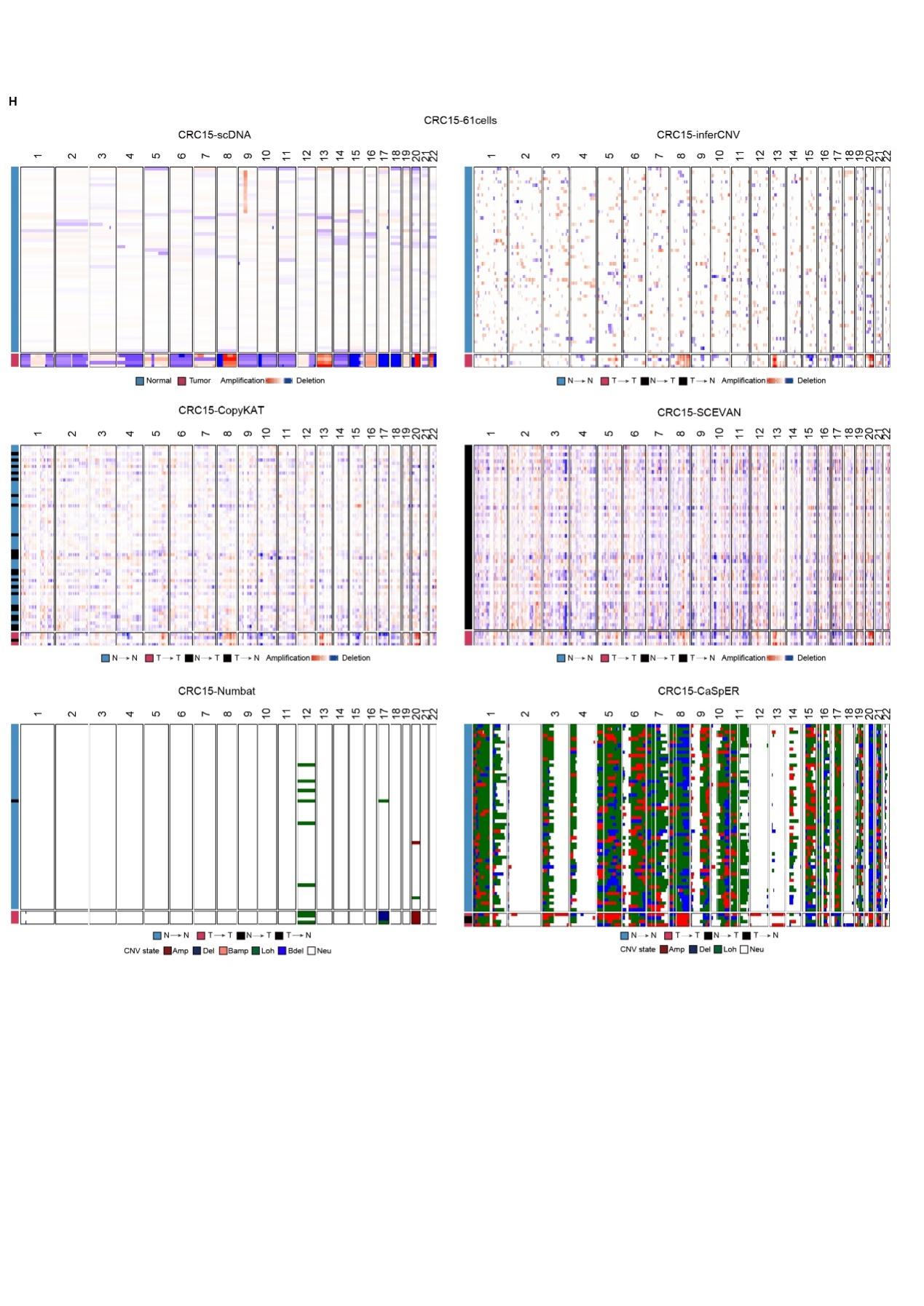

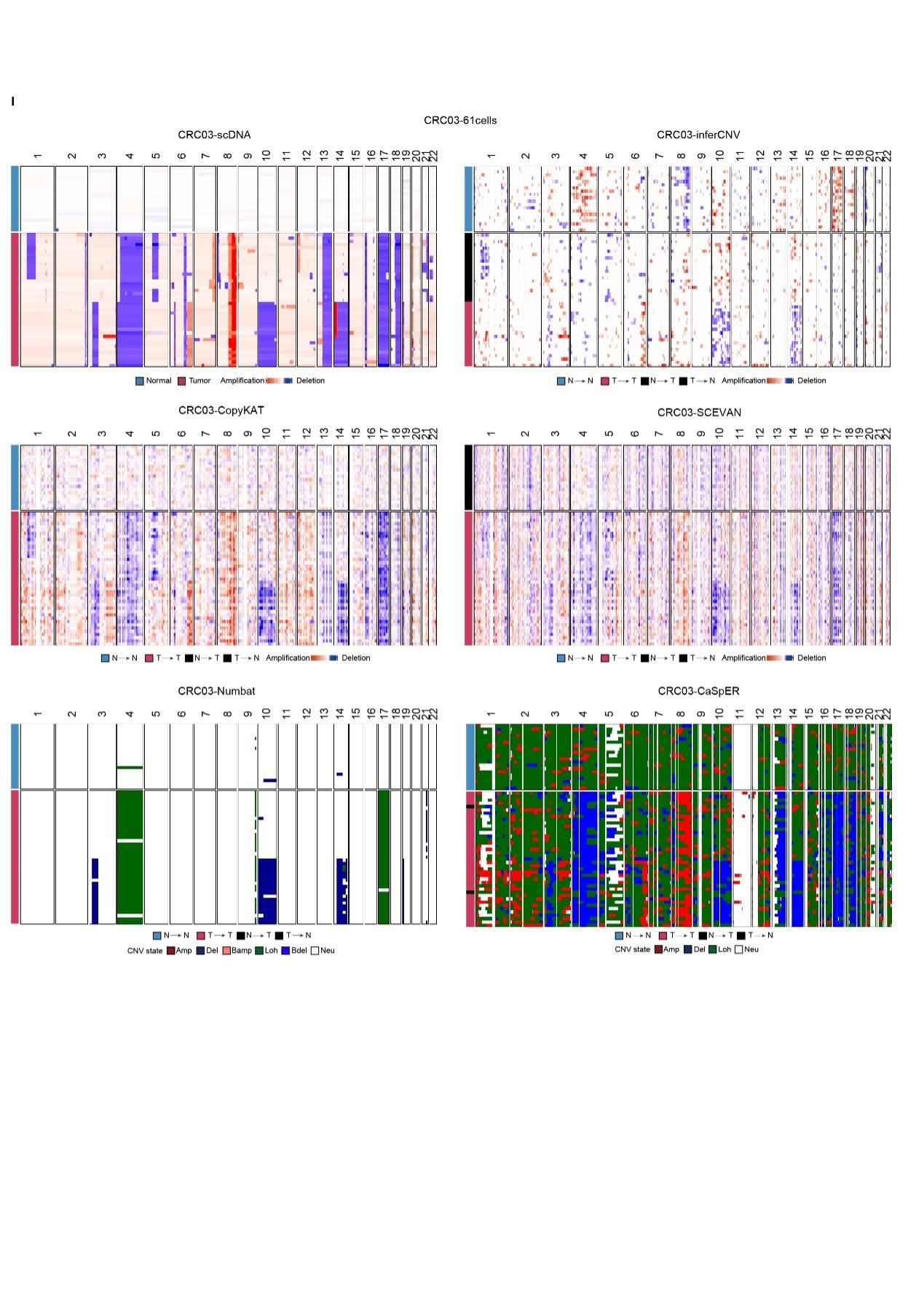

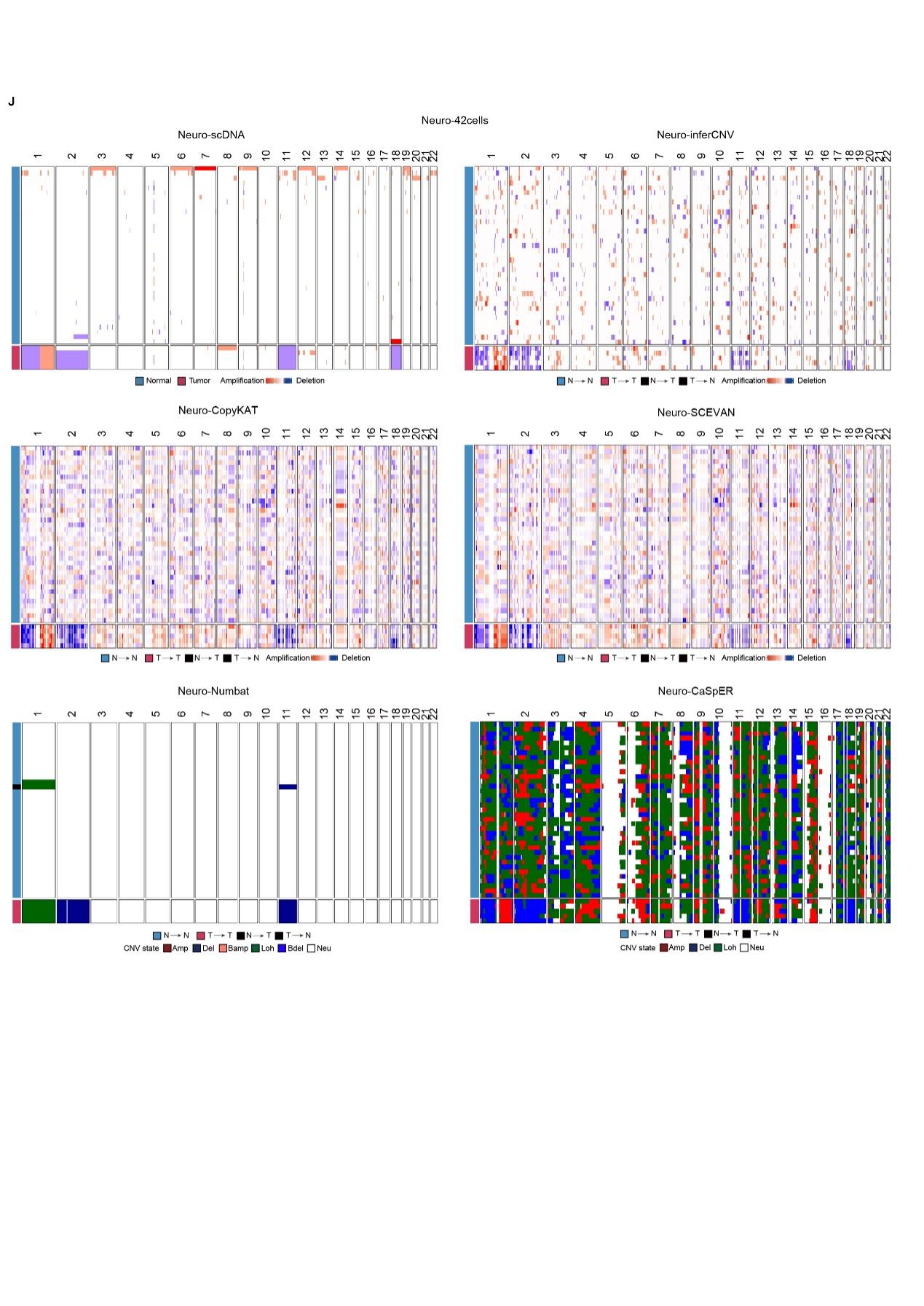

**Supplementary FigureS1: Heatmap displaying the CNAs profiles of glioma, CRC and neuroendocrine tumor patients**

(A-J) Heatmap showing the CNAs profiles obtained from ground truth scDNA-seq, inferred by inferCNV, CopyKAT, SCEVAN, Numbat, and CaSpER in glioma (A), CRC13 (B), CRC11 (C), CRC12 (D), CRC16 (E), CRC02 (F), CRC22 (G), CRC15 (H), CRC03 (I), Neuro(J).

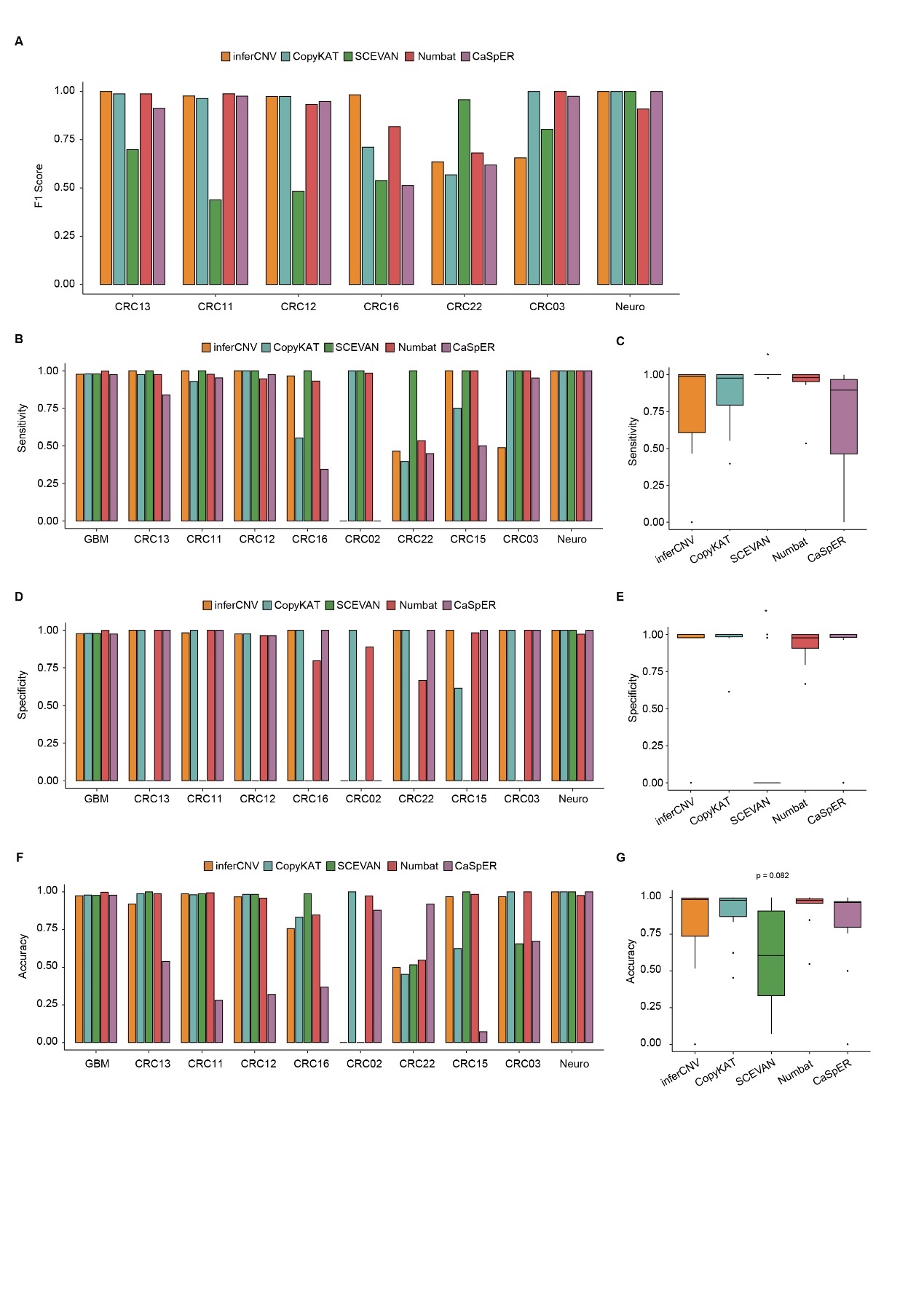

**Supplementary FigureS2: Evaluation metrics for tumor and normal cells classification**

A: The F1 scores for tumor and normal cell classification in CRC13, CRC11, CRC12, CRC16, CRC22, CRC03 and Neuro.

B&D&F: Bar plots showing Sensitivity (B), Specificity(D) and Accuracy(F) for tumor and normal cell classification for various tools.

C&E&G: Boxplots showing Sensitivity (C), Specificity(E) and Accuracy(G) for tumor and normal cell classification for various tools. And p values in (C, E and G) calculated using Kruskal-Wallis test. NS: p > 0.05, *: p < 0.01, **: p < 0.001, ***: p < 0.0001

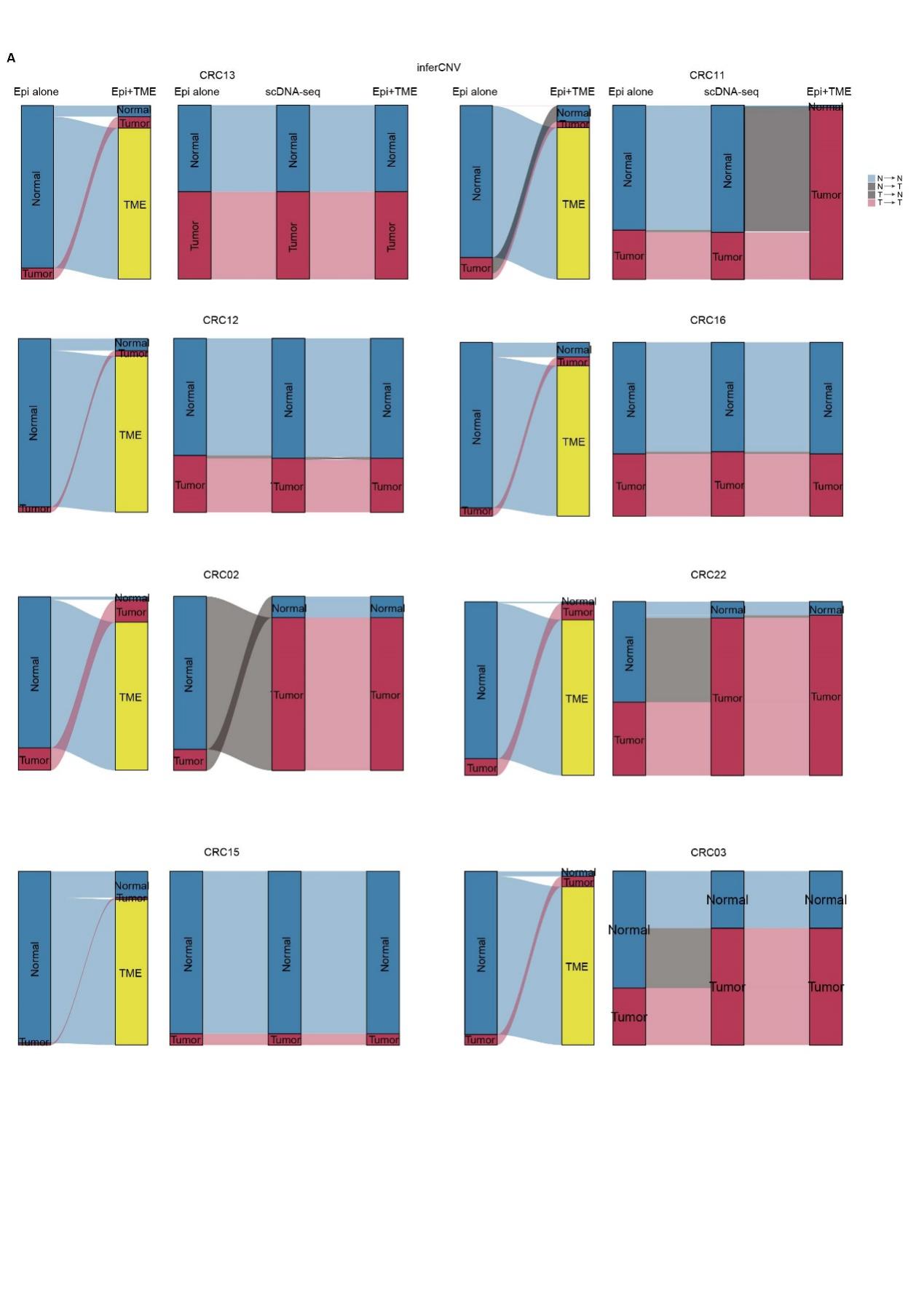

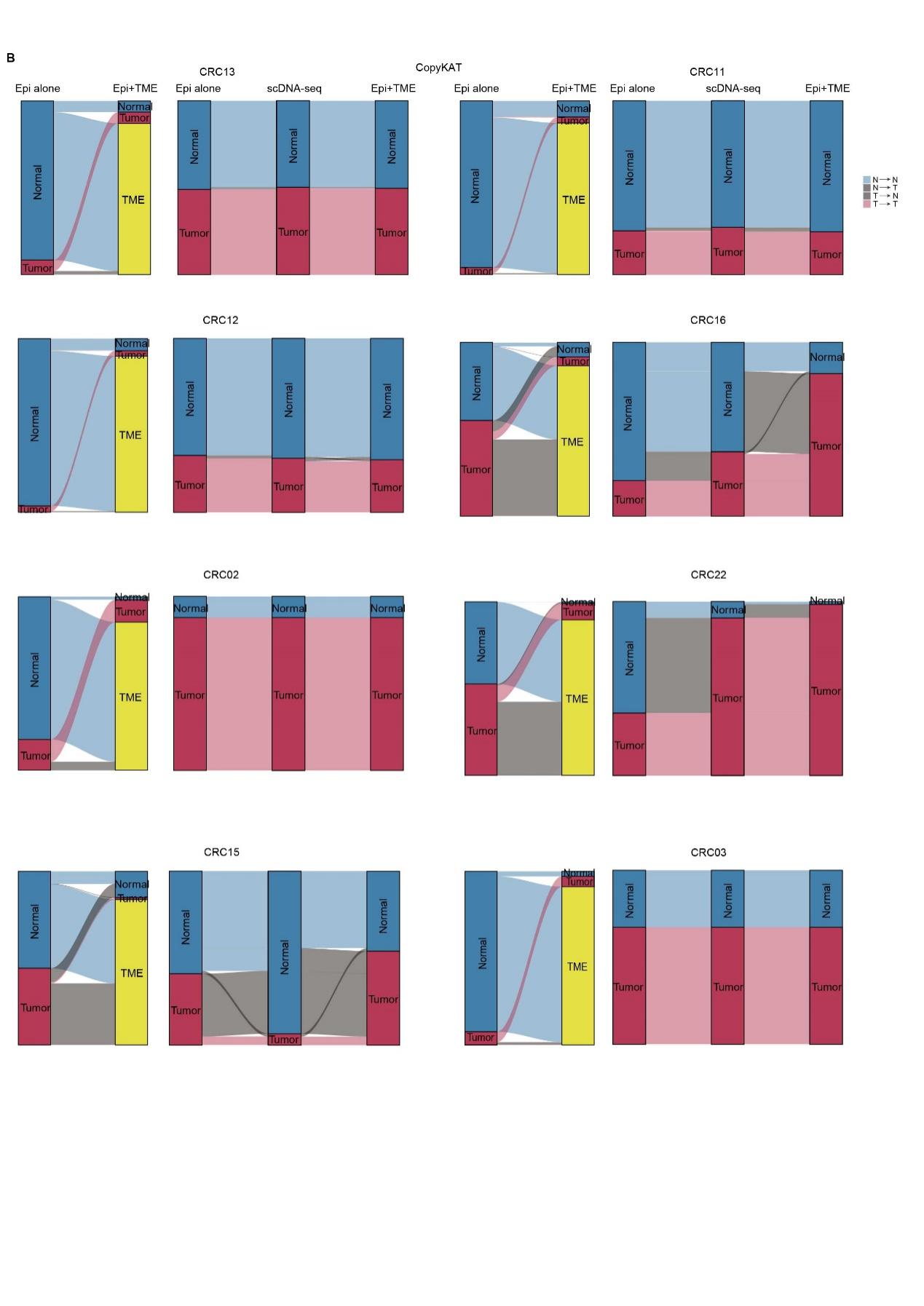

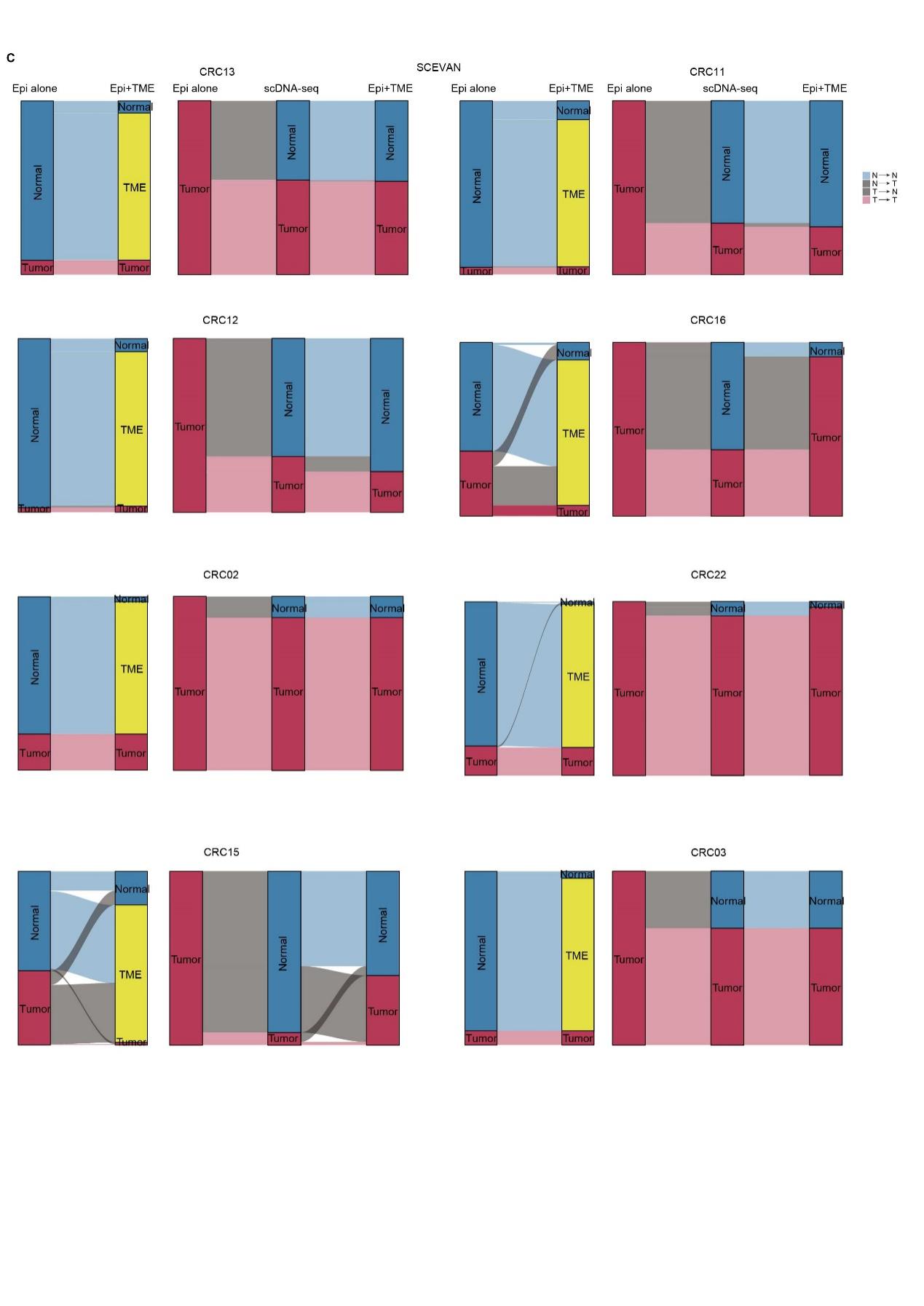

**Supplementary FigureS3: Sankey diagrams illustrating the classification of tumor and normal cells under epithelial alone or epithelial inclusion TME cells from CRC cohort**

A-C: Sankey diagrams showing the classification predicted by inferCNV (A), CopyKAT (B) and SCEVAN (C). For each sub-figure, the left panel shows the overall tumor vs. normal cells classification, and the right panel exhibits the tumor vs. normal cells classification of epithelial cells under two conditions: epithelial alone and epithelial including TME cells.

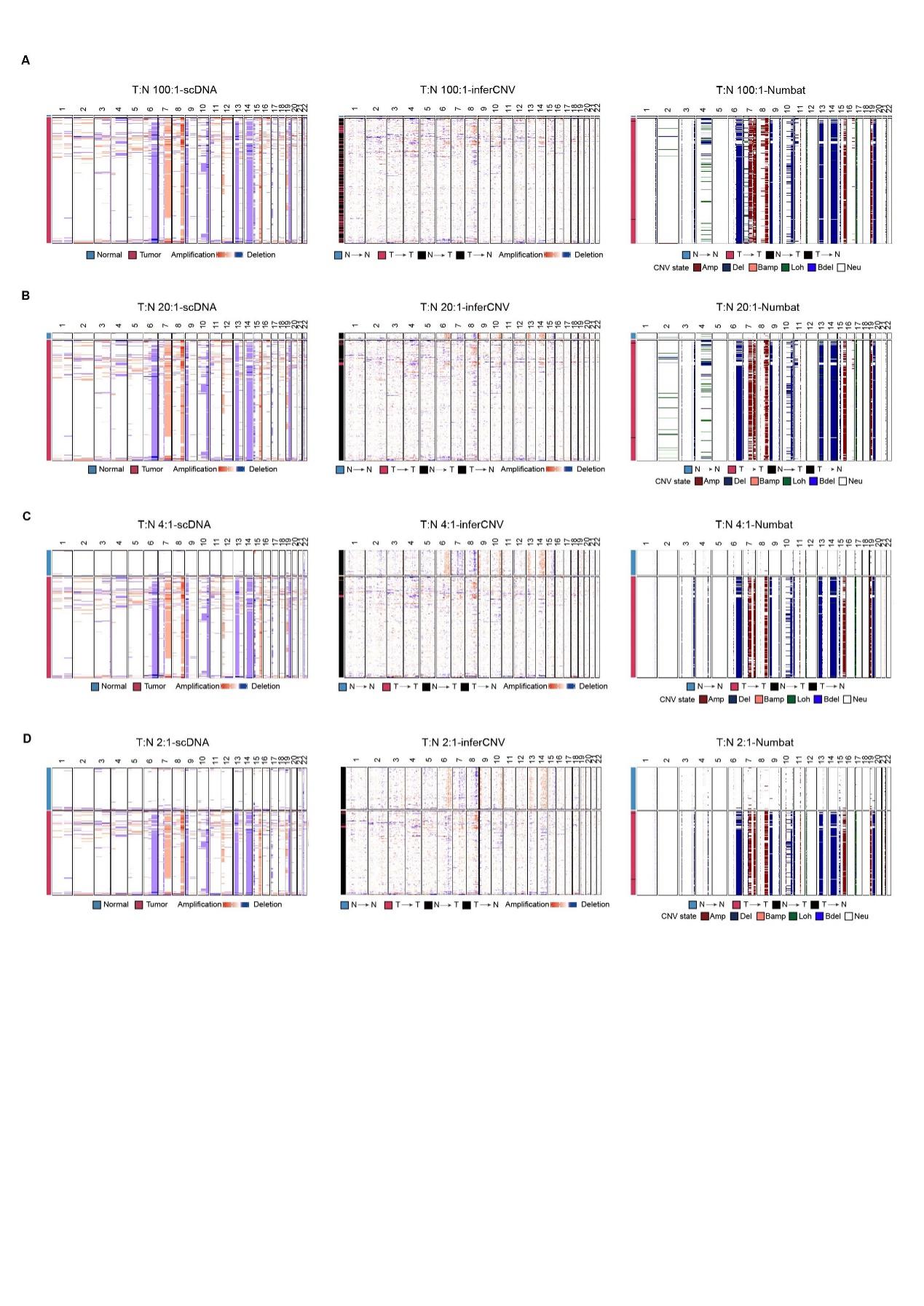

**Supplementary FigureS4: Heatmap displaying the CNAs profiles of downsampling data**

A-D: CNAs profiles from scDNA, inferCNV and Numbat under the ratio of sampling tumor and normal cells number is 100:1 (A), 20:1 (B), 4:1 (C) and 2:1 (D).

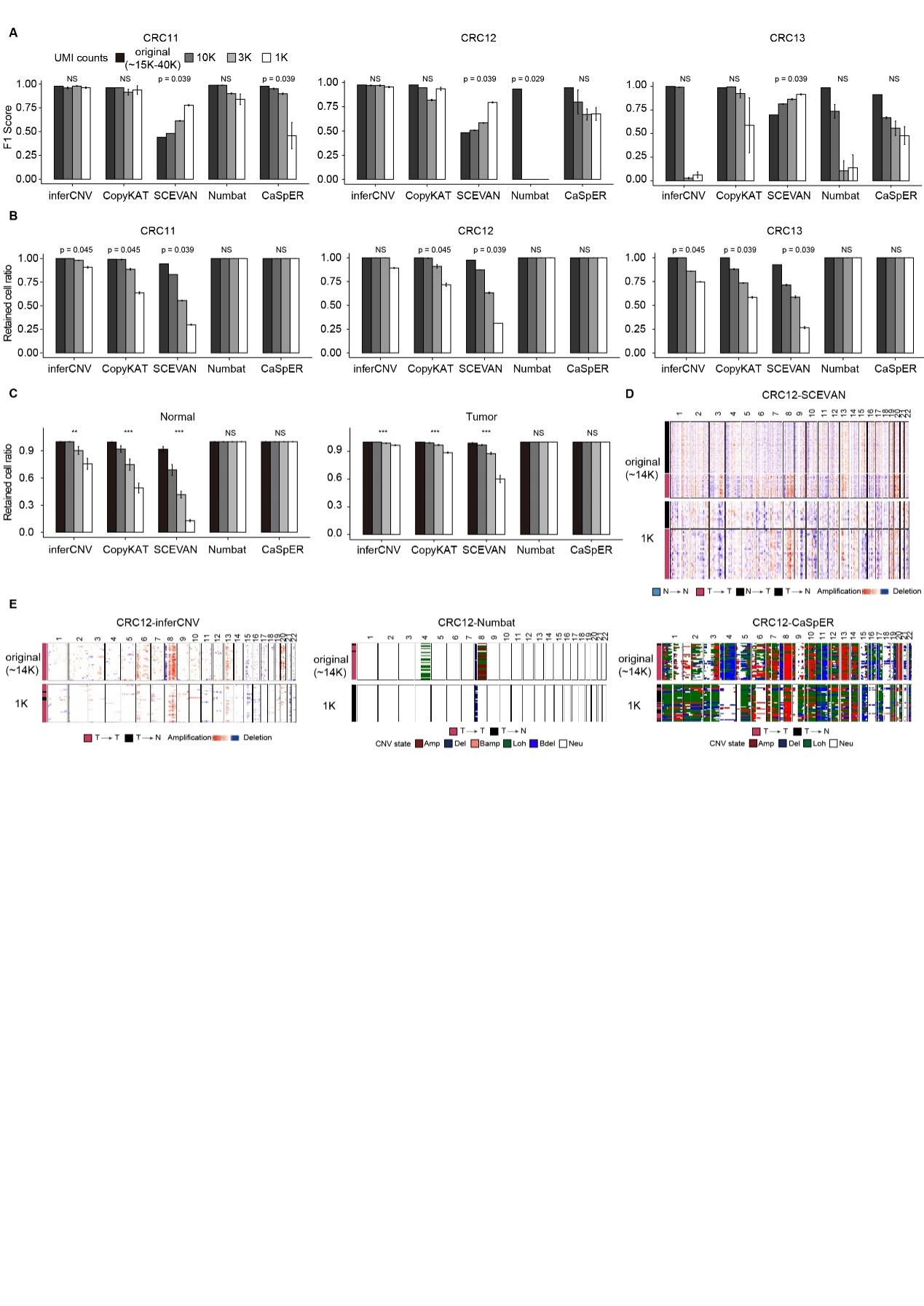

**Supplementary Figure S5: Benchmark of tumor and normal cells classification** **under different sequencing depths**

A: F1 scores for tumor and normal cell classification under different sequencing depths in CRC11, CRC12 and CRC13.

B: The ratios of cells passing quality control at different sequencing depths in CRC11, CRC12 and CRC13.

C: The ratios of Normal and Tumor cells passing quality control at different sequencing depths in the combined CRC11, CRC12 and CRC13.

D: The heatmap represents the tumor cells and normal cells CNAs profiles of SCEVAN affected by sequencing depths. The upper heatmap represents CNAs profiles from original scRNA-seq, and the bottom heatmap exhibits CNAs profiles from down-sampled low-sequencing depth (1K). The number of normal cells filtered out due to low sequencing depth decreased much more rapidly compared to tumor cells. (refer to C panel of this figure) which led to much lower normal cell percentage in 1k versus original sequencing depth.

E: The heatmaps represent the tumor cells CNAs profiles of inferCNV, Numbat and CaSpER affected by sequencing depths. The upper heatmaps represent CNAs profiles from original scRNA-seq, and the bottom heatmaps show CNAs profiles from down-sampled low-sequencing depth (1K).

And p values in (A-C) calculated using Kruskal-Wallis test. NS: p > 0.05, *: p < 0.01, **: p < 0.001, ***: p < 0.0001

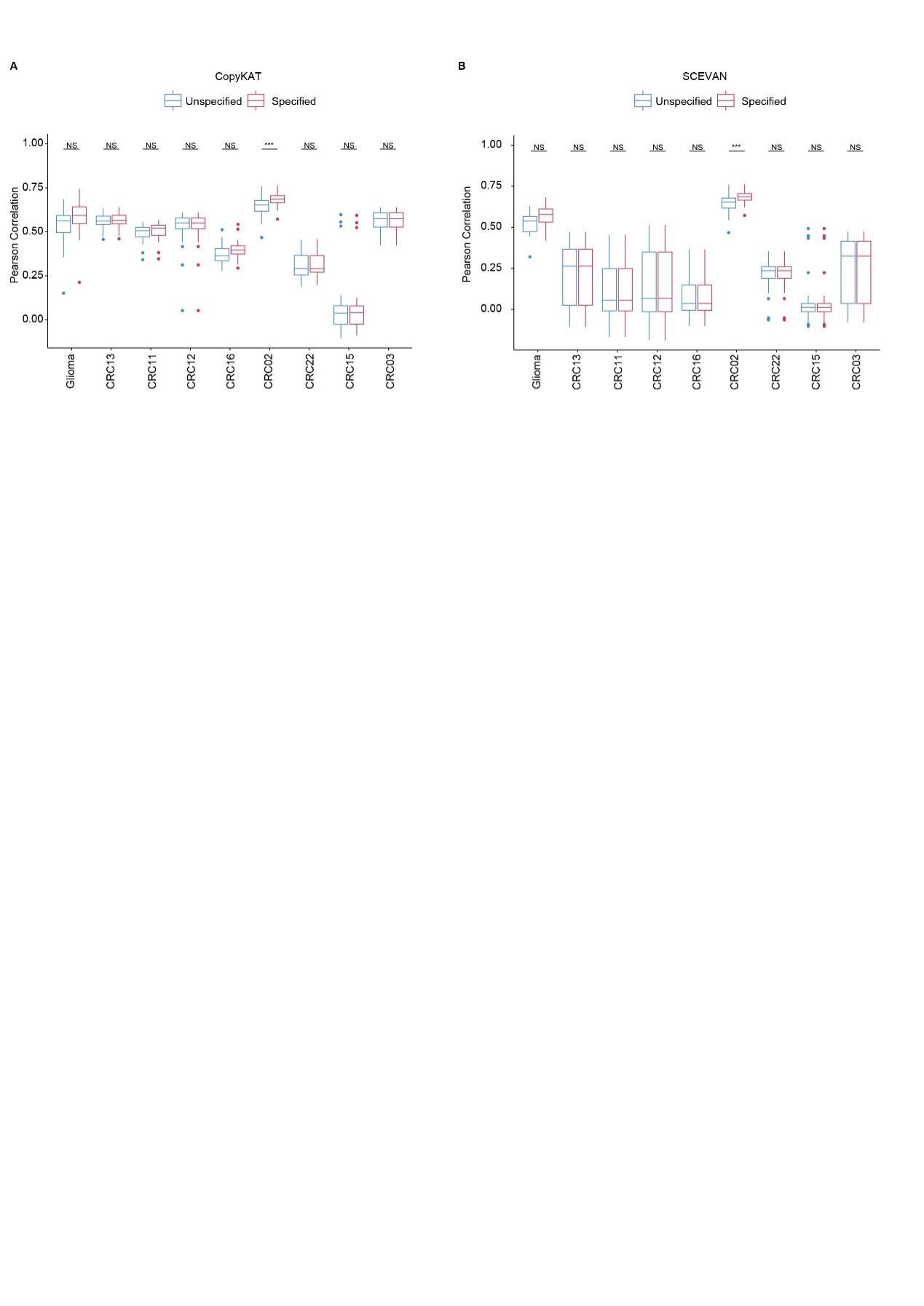

**Supplementary FigureS6: Pearson correlation of CNAs between scDNA-seq and scRNA-seq under unspecified and specified reference cells**

A-B: Boxplots showing the Pearson correlation between inferred CNAs from scRNA-seq and the ground truth from scDNA-seq for each patient under unspecified and specified reference cells with CopyKAT (A) and SCEVAN (B).

p-values in (A-B) calculated using unpaired two-tailed Wilcoxon rank-sum test, corrected using the Benjamini–Hochberg method for (A-B). NS: p > 0.05, *: p < 0.01, **: p < 0.001, ***: p < 0.0001

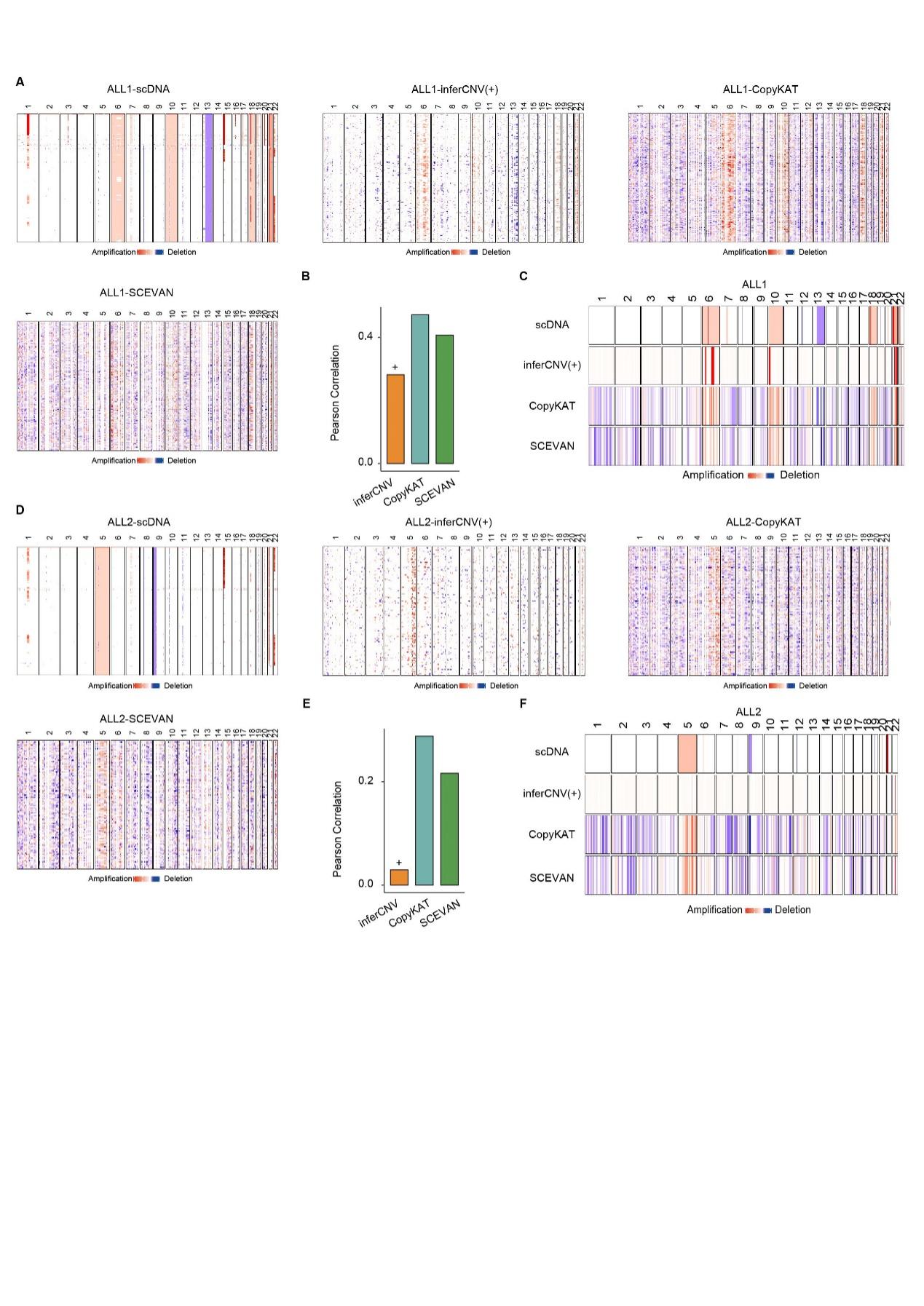

**Supplementary FigureS7: CNAs profiles for acute lymphoblastic leukemia**

A&D: Heatmap showing the CNAs profiles of blasts tumor cells from scDNA, inferCNV, CopyKAT and SCEVAN for ALL1 (A) and ALL2 (D) patients.

B&E: Bar plots showing Pearson correlation of tumor cells between inferred CNAs from scRNA-seq and the ground truth from scDNA-seq for ALL1 (B) and ALL2 (E) patients across inferCNV, CopyKAT and SCEVAN.

C&F: Heatmap showing the median CNAs profiles of blasts tumor cells from ground truth scDNA-seq, inferCNV, CopyKAT and SCEVAN for ALL1 (C) and ALL2 (F) patients.

"+" indicates tumor cell CNAs profiles estimated when both tumor and TME cells are inputted into inferCNV.

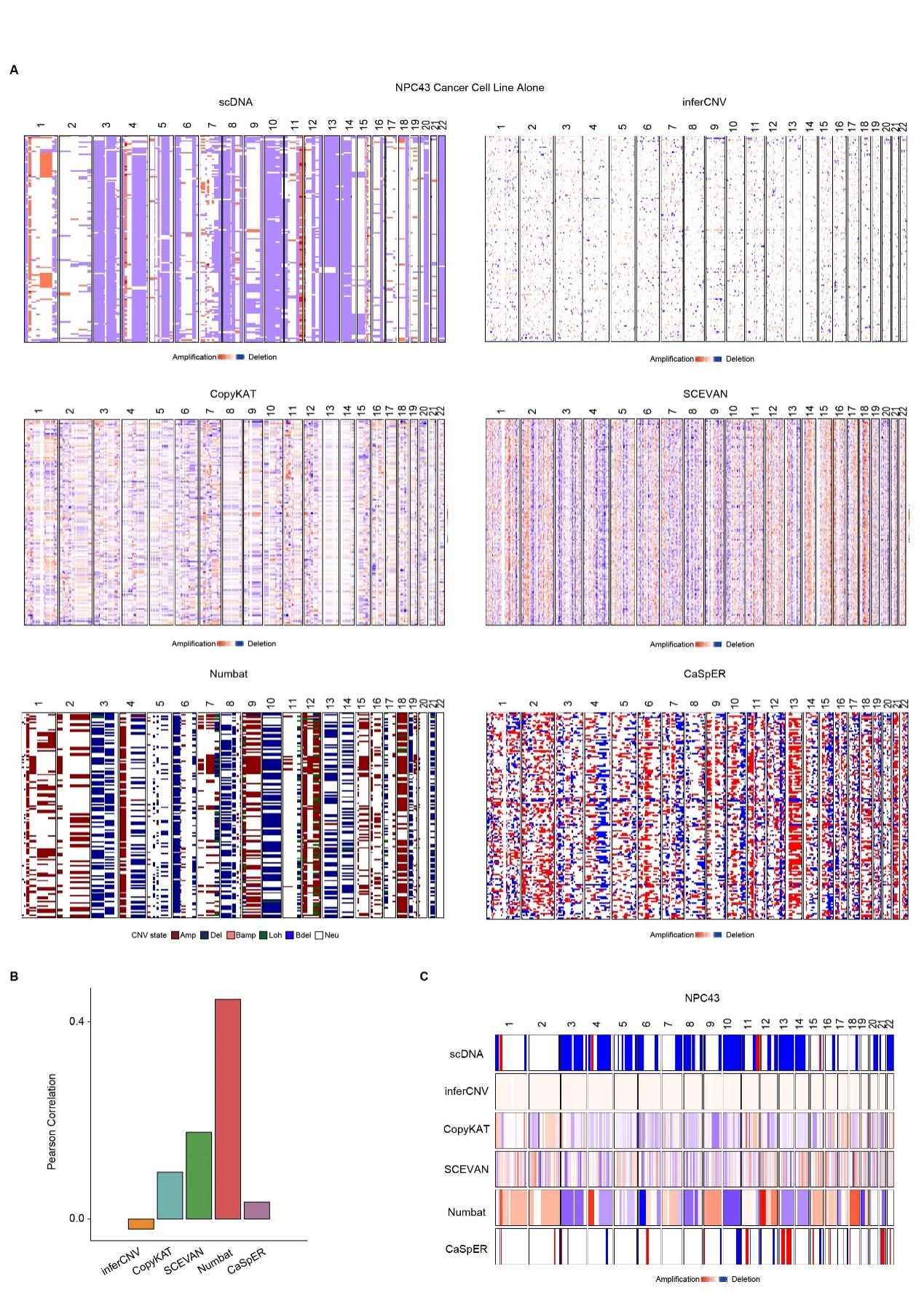

**Supplementary FigureS8: CNAs profiles for NPC43 cell line without setting reference**

A: Heatmap showing the CNAs profiles of blasts tumor cells from scDNA, inferCNV, CopyKAT, SCEVAN, Numbat and CaSpER.

B: Bar plots showing the CNAs profiles Pearson correlation of NPC43 cell line between the scDNA-seq and inferred by inferCNV, CopyKAT, SCEVAN, Numbat and CaSpER.

C: Heatmap showing the CNAs profiles of NPC43 cell line from ground truth scDNA-seq, inferCNV, CopyKAT, SCEVAN, Numbat and CaSpER.

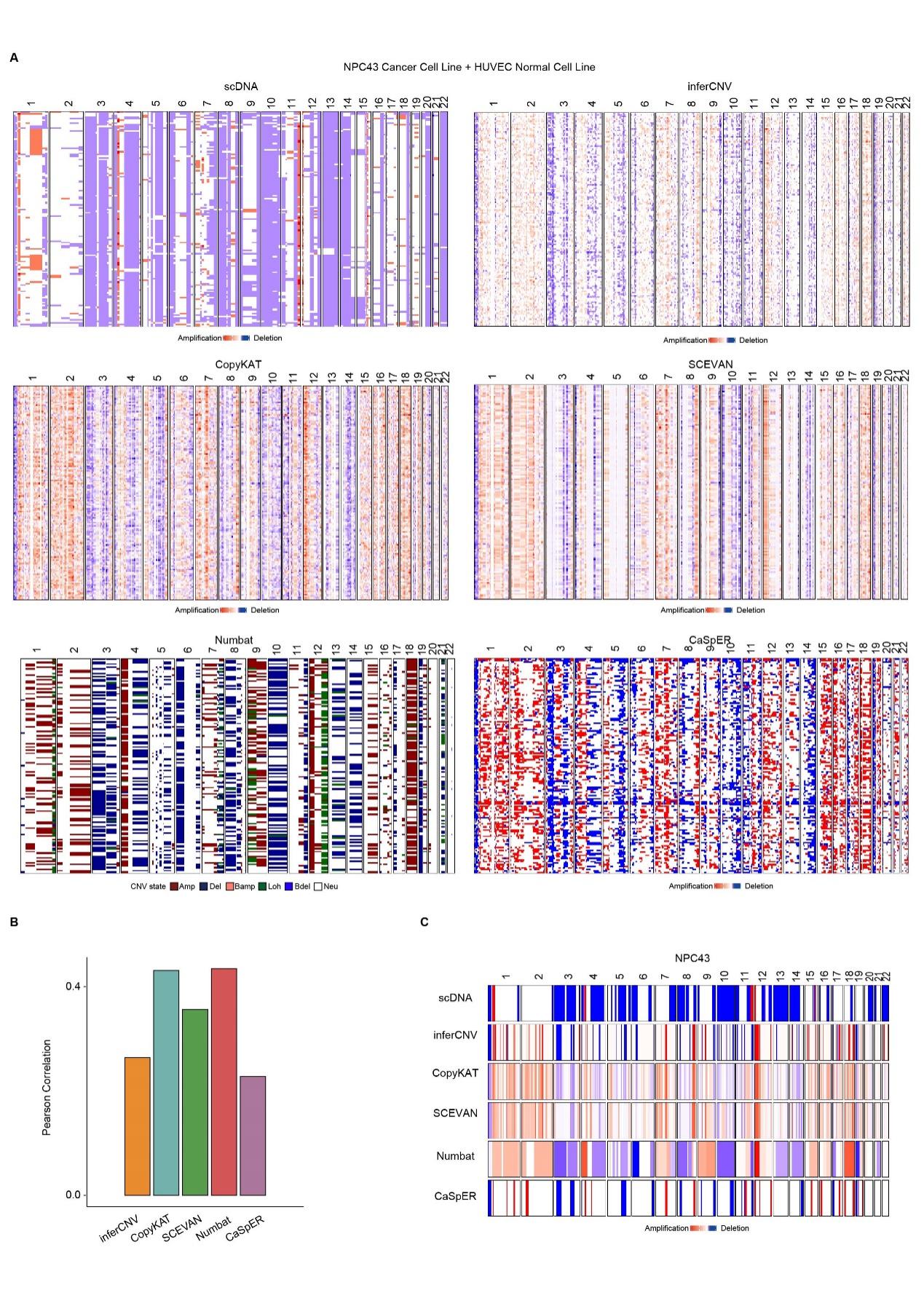

**Supplementary FigureS9: CNAs profiles for NPC43 cell line with HUEVC cells as normal reference**

A: Heatmap showing the CNAs profiles of blasts tumor cells from scDNA, inferCNV, CopyKAT, SCEVAN, Numbat and CaSpER.

B: Bar plots showing the CNAs profiles Pearson correlation of NPC43 cell line between the scDNA-seq and inferred by inferCNV, CopyKAT, SCEVAN, Numbat and CaSpER.

C: Heatmap showing the median CNAs profiles of NPC43 cell line from ground truth scDNA-seq, inferCNV, CopyKAT, SCEVAN, Numbat and CaSpER.

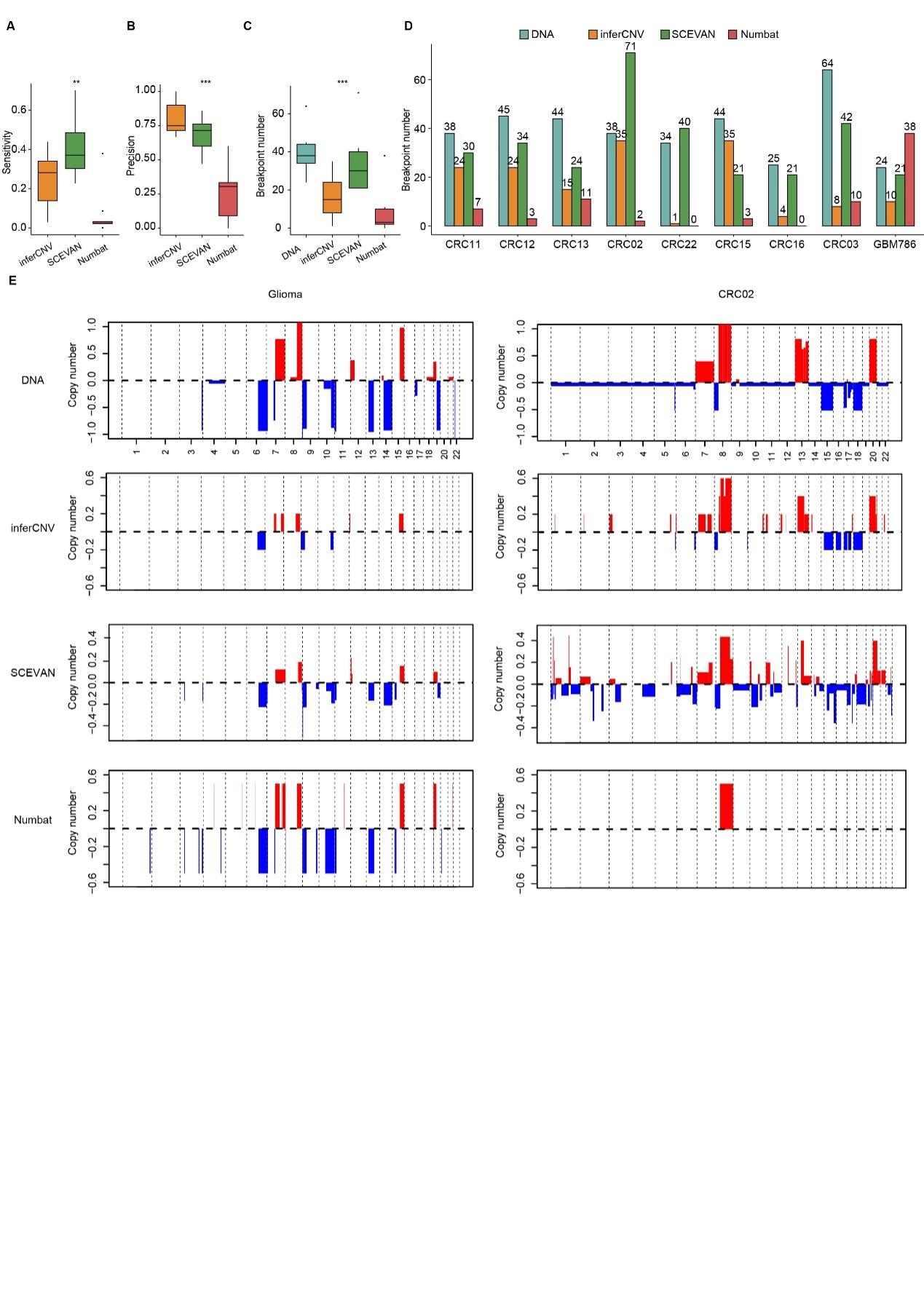

**Supplementary Figure S10:** **Benchmark of breakpoints in inferCNV, SCEVAN and Numbat**

A: Boxplot of Sensitivity for breakpoints detection in inferCNV, SCEVAN and Numbat.

B: Boxplot of Precision for breakpoints detection in inferCNV, SCEVAN and Numbat

C: Boxplot of the number of breakpoints detected in inferCNV, SCEVAN and Numbat

D: A bar plot illustrates the number of breakpoints detected in each tumor sample

E: genome wide CNAs (Breakpoints) from glioma and CRC02 samples

P values in (A-C) calculated using Kruskal-Wallis test. NS: p > 0.05, *: p < 0.01, **: p < 0.001, ***: p < 0.0001

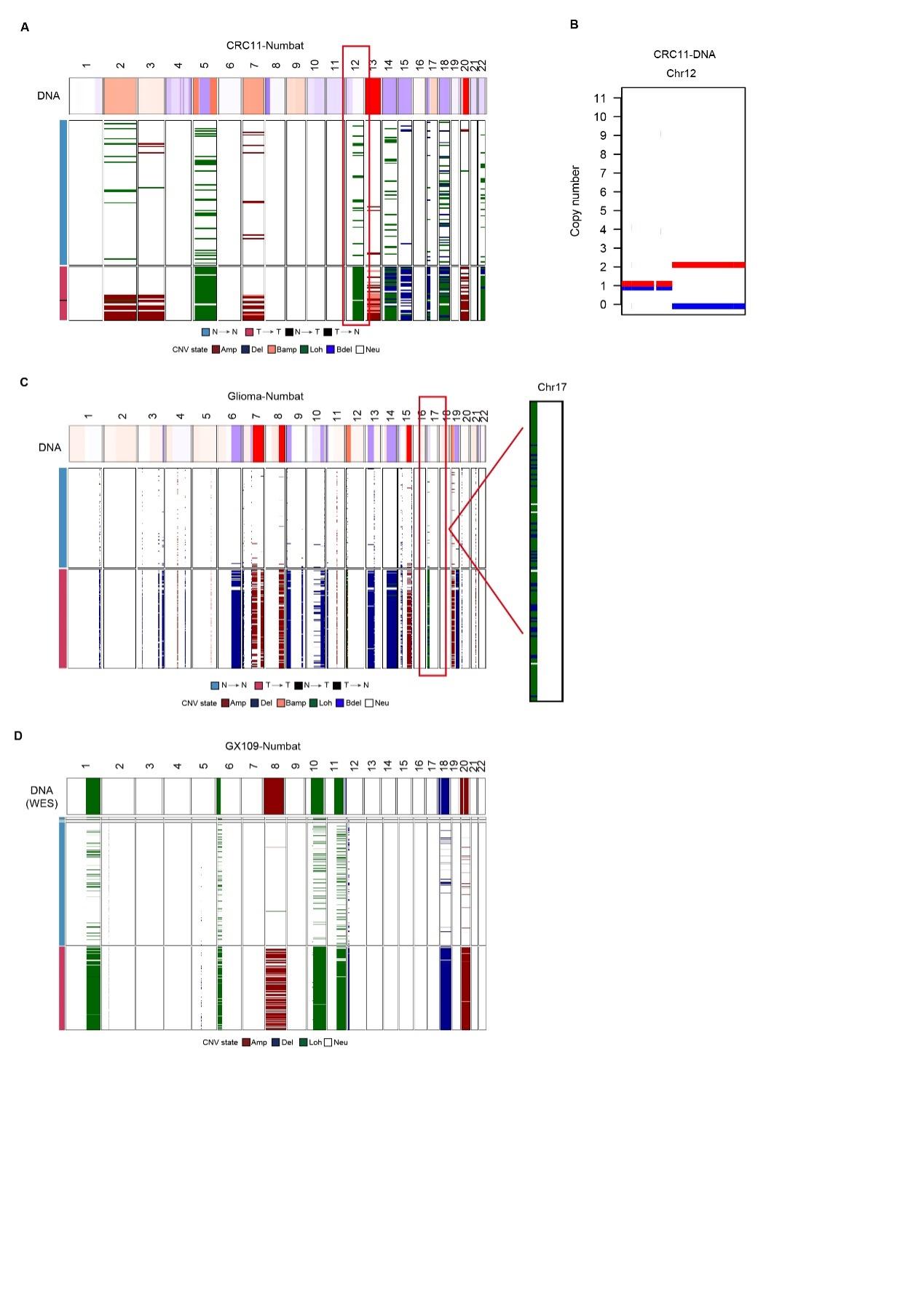

**Supplementary Figure S11: validation of Numbat detected LOH events by paired DNA-Seq.**

A: Heatmap showing the CNAs profiles of tumor/normal cells from Numbat in CRC11, with the red box highlighting the chr12 LOH.

B: Sequenza CNV AB allele copy number from CRC11 validated the chr12 LOH event.

C: Heatmap showing the CNAs profiles of tumor/normal cells from Numbat in Glioma with a zoomed-in view of LOH events on 17p shown on the right.

D: Heatmap showing the CNAs profiles of tumor/normal cells from Numbat in GX109.

The paired DNA SCNA profiles for tumors were shown on the top panel for A,C,D.

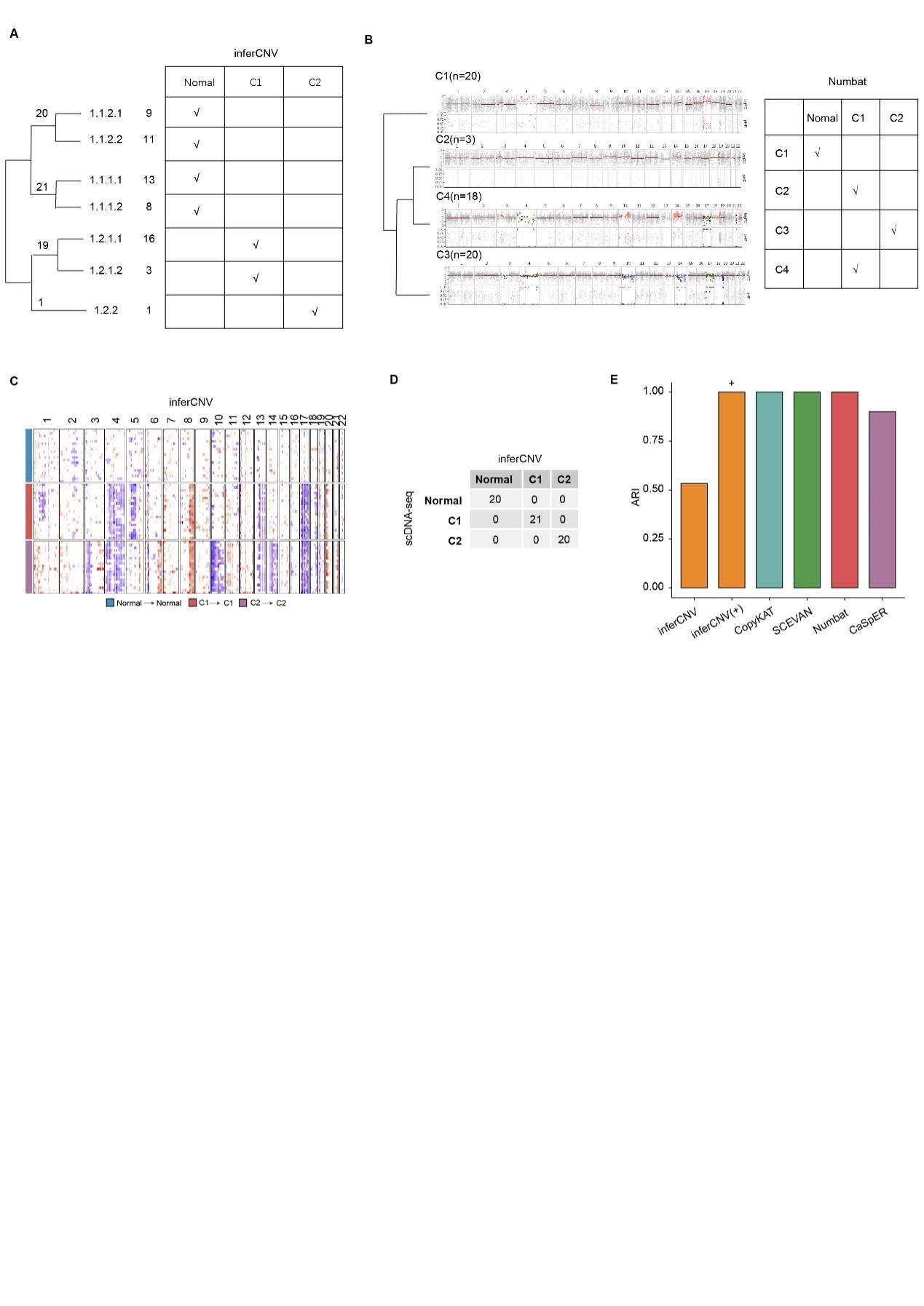

**Supplementary Figure S12: inferCNV CRC03 tumor subclone structure inference with TME cells included**

A: Cartoon illustrates how multiple subclones from inferCNV were combined based on CNAs profile similarity into one normal and two tumor subclones

B: Cartoon illustrates how multiple subclones from Numbat were combined based on CNAs profile similarity into one normal and two tumor subclones

C: Heatmap displaying the CNAs profiles of CRC03 subclones after inclusion of TME cells

D: Tables presenting the cell numbers corresponding to ground truth subclone labels from scDNA and predicted subclone labels of CRC03 after including TME cells.

E: Bar plot showing the performance evaluation ARI of CRC03 tumor subclone inferred by different tools.

"+" indicated tumor subclone estimated when TME cells were included.

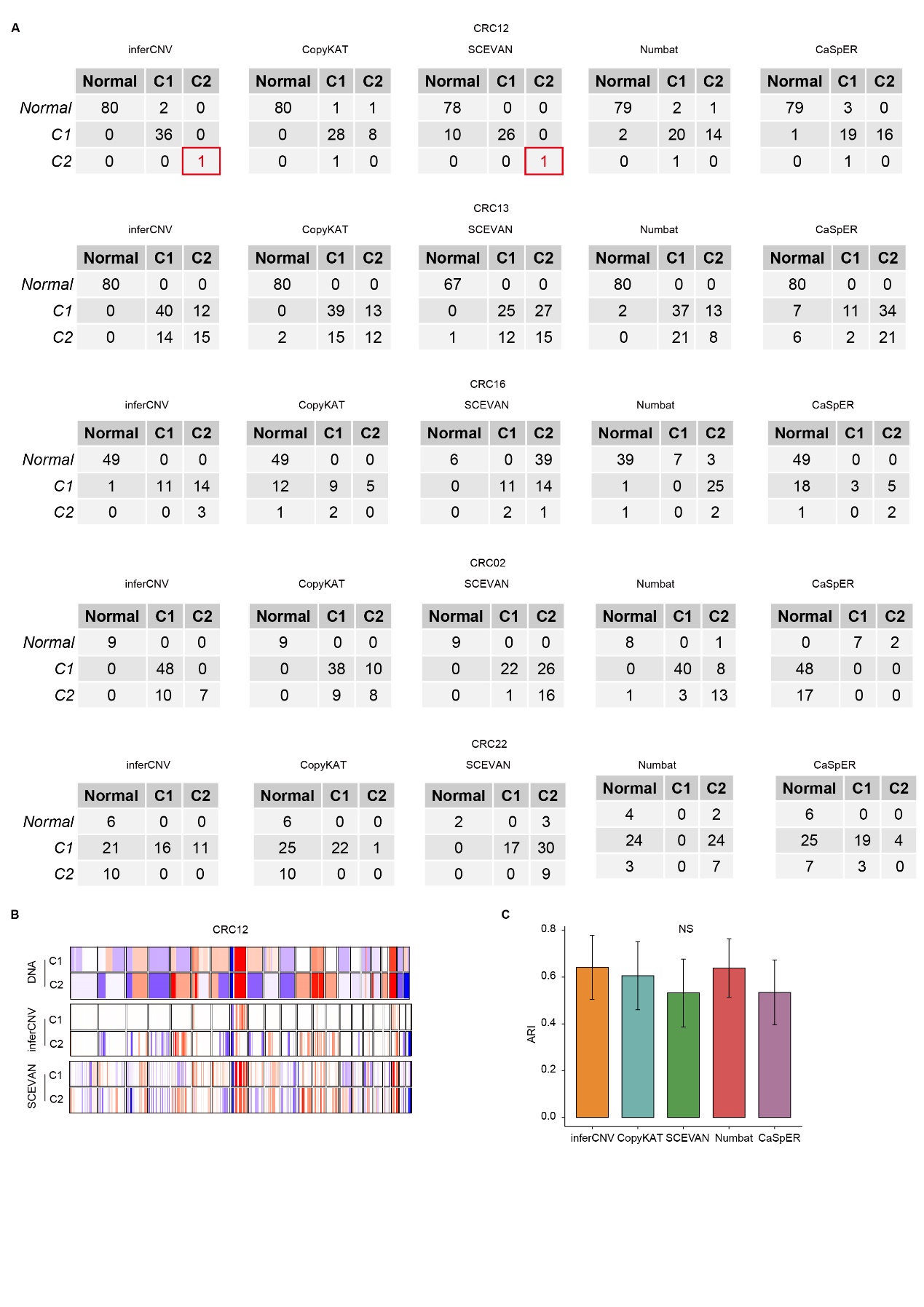

**Supplementary FigureS13:** Tumor clonal evolution inference

A: Tables exhibit cell numbers associated with subclone labels from scDNA and the predicted subclone labels from scRNA for CRC12, CRC13, CRC16, CRC02 and CRC22. The red color “1” indicates an extreme case where one tumor subclone is only consisted of one single, which was correctly detected by inferCNV and SCEVAN.

B: Heatmaps comparing the CNAs profiles of two subclones in CRC12 from scDNA-seq, inferCNV and SCEVAN, with C2 subclone contained only one cell.

C: Bar plots show the Adjusted Rand Index (ARI) for between scDNA-seq and inferred subclones

And P value in C calculated using Kruskal-Wallis test, NS represents statistically nonsignificant >0.05, P value. NS: p > 0.05, *: p < 0.01, **: p < 0.001, ***: p < 0.0001

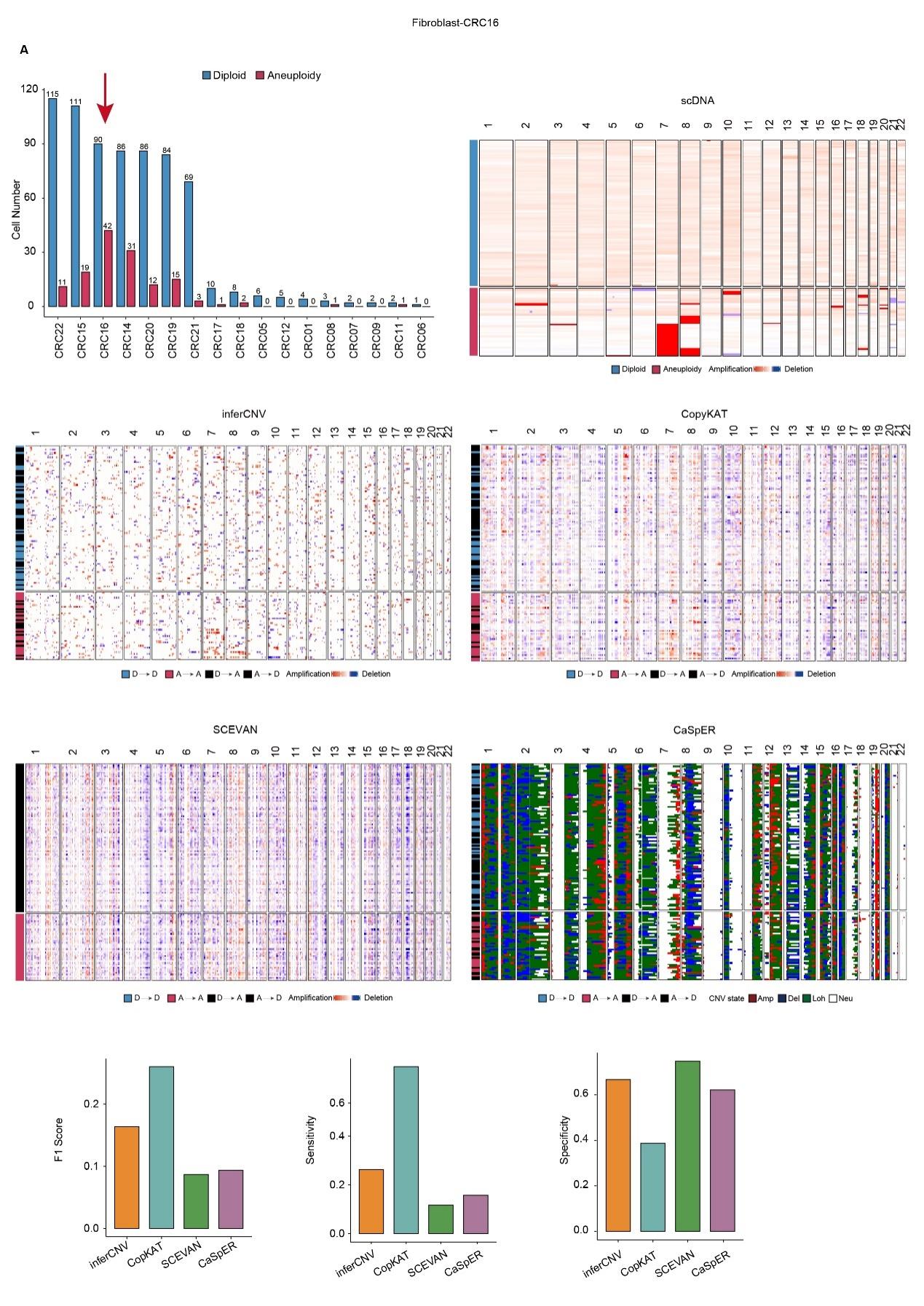

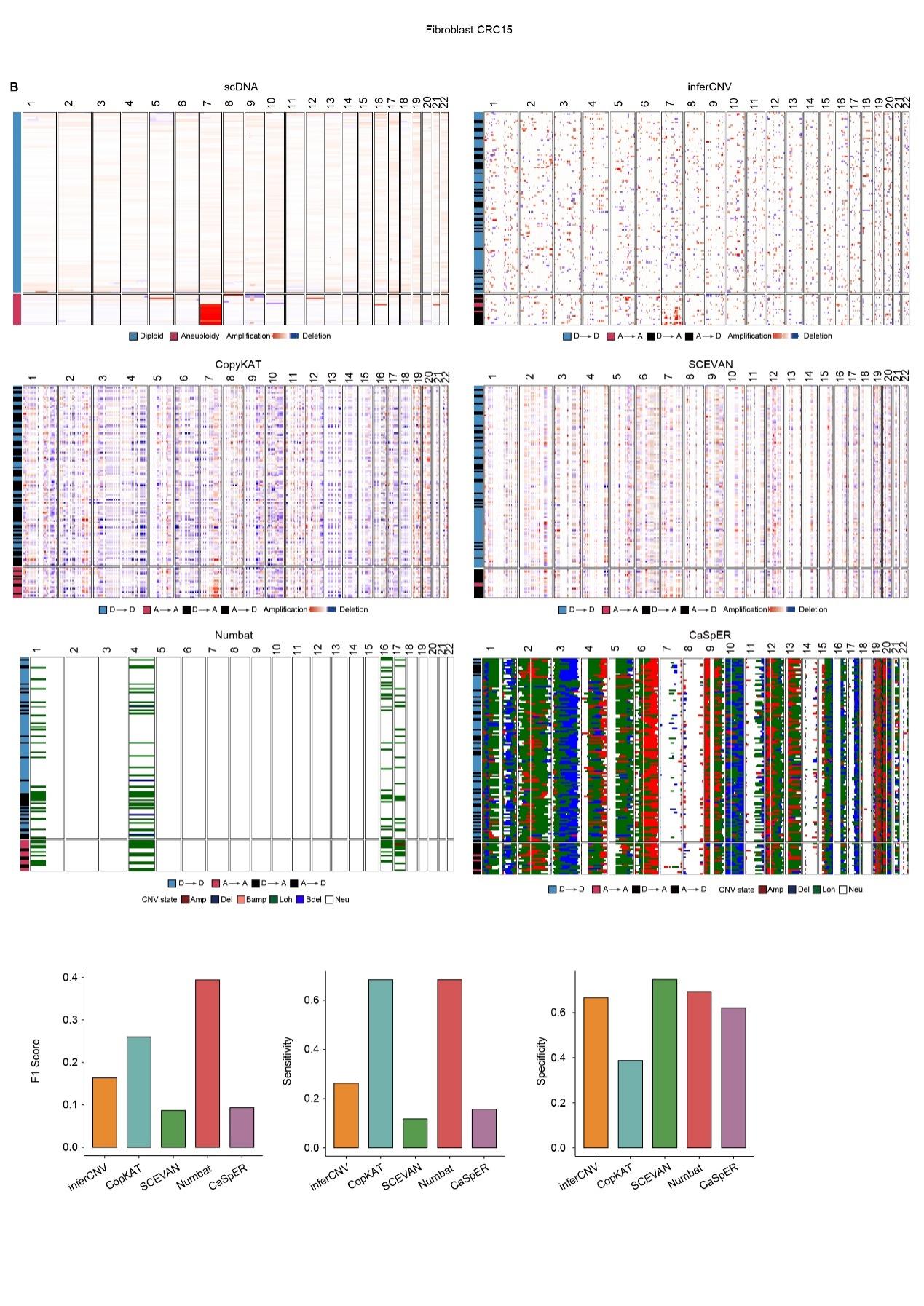

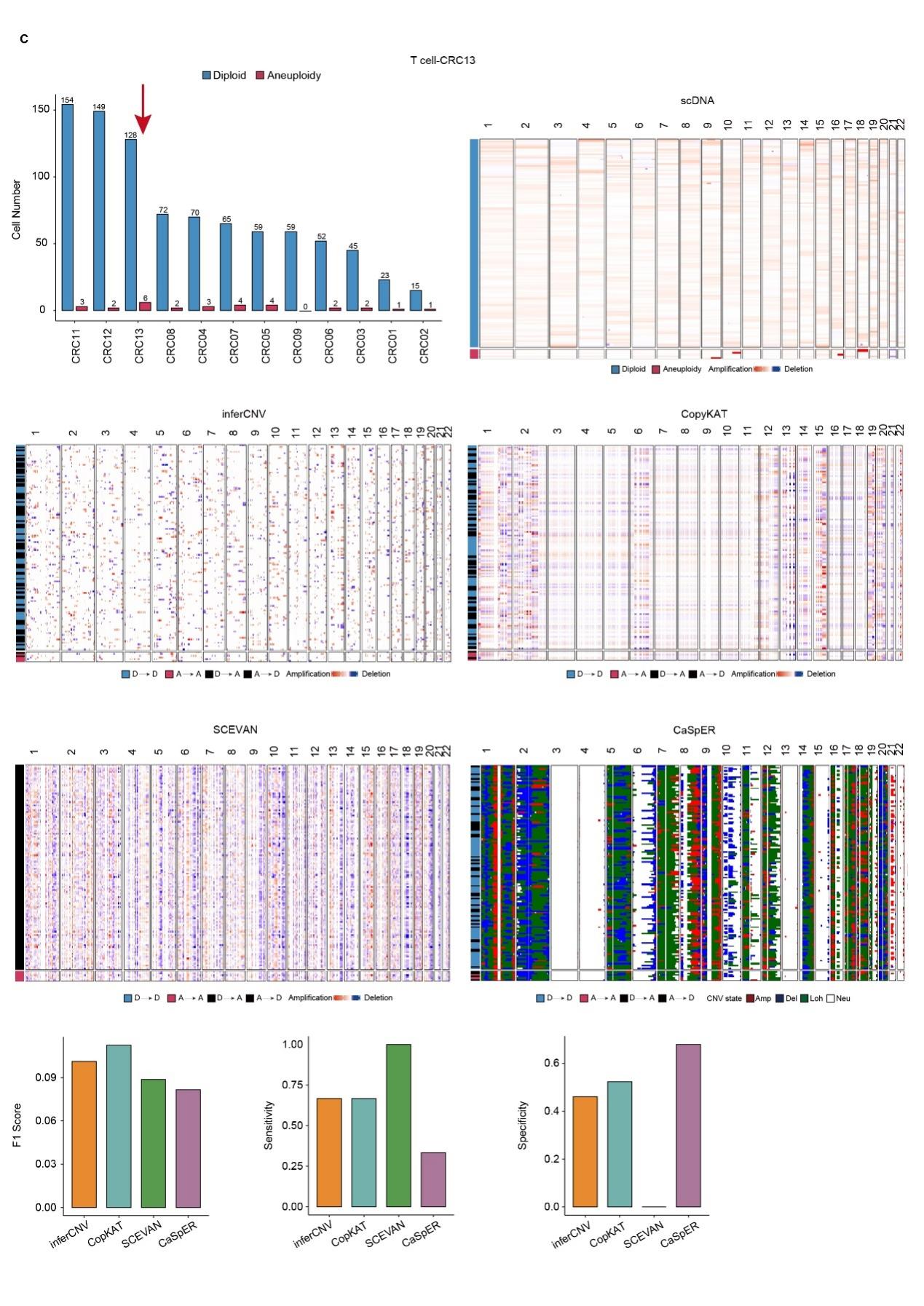

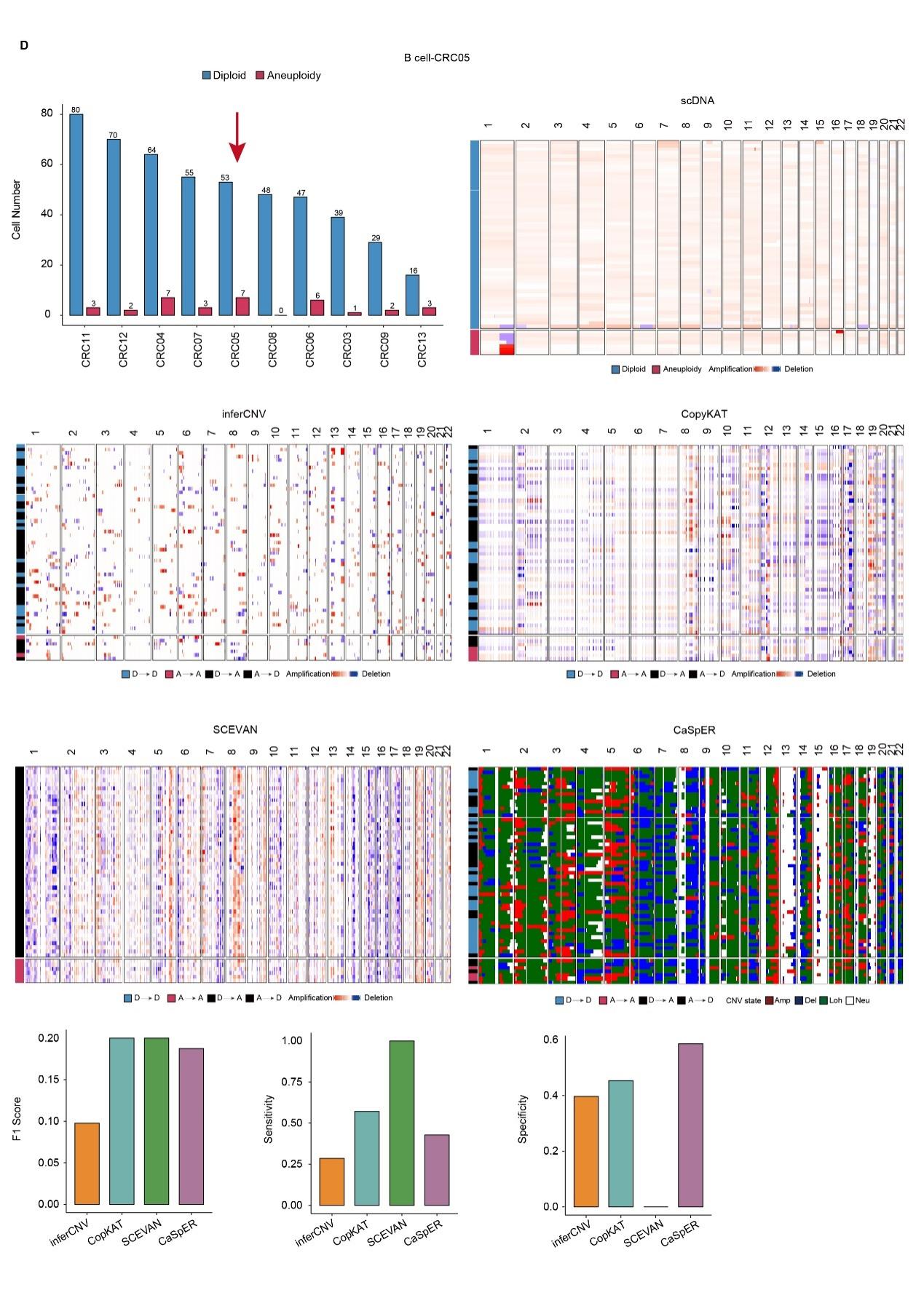

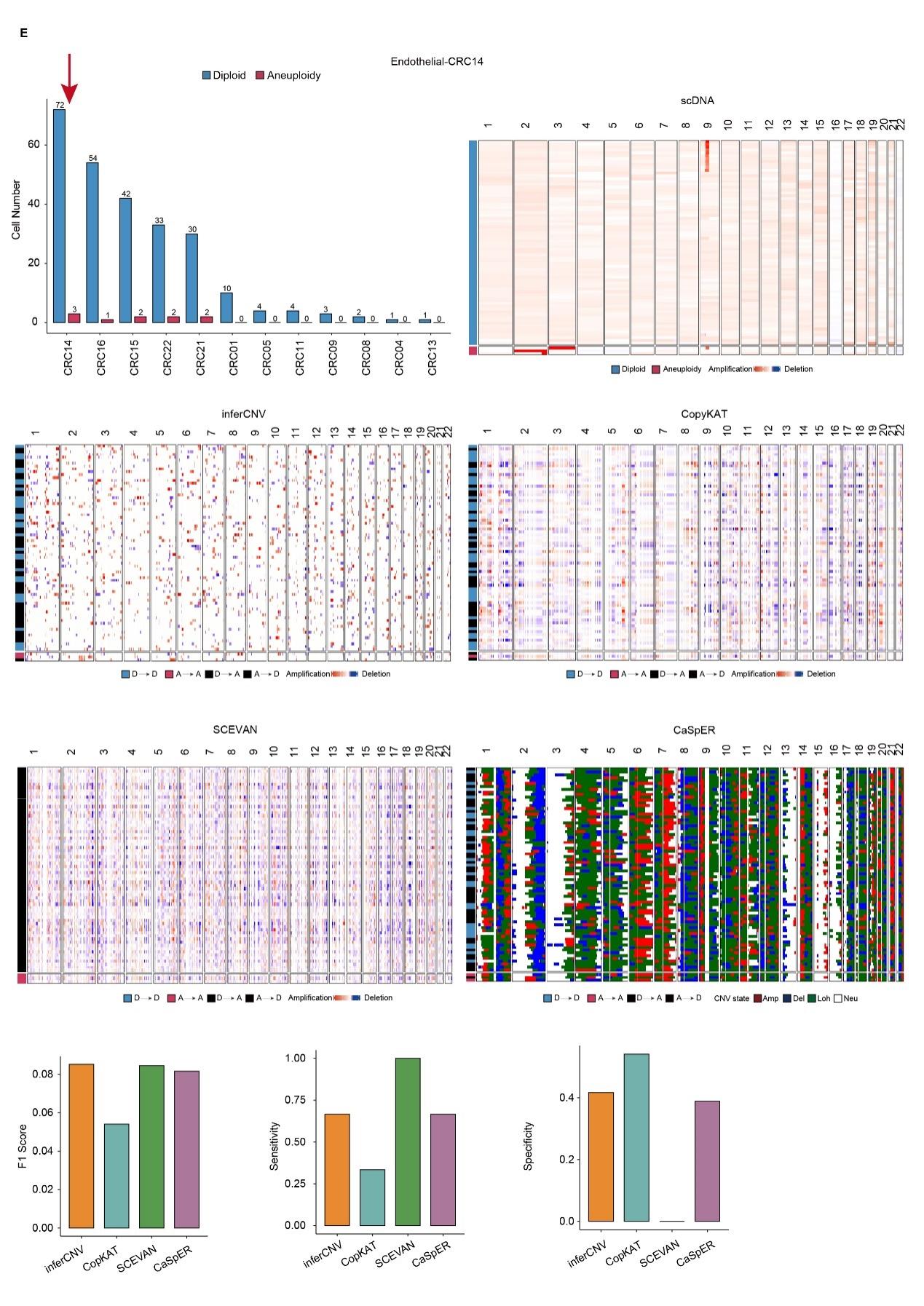

**Supplementary FigureS14: Benchmark of aneuploidy inference in normal cell**

A-E: The counts of autosomal aneuploidy and euploid cells for each patient. Heatmap showing the CNAs profiles of selected patients indicated by the arrow obtained from ground truth scDNA-seq, inferred from inferCNV, CopyKAT, SCEVAN, Numbat and CaSpER. F1 scores, Sensitivity and Specificity for aneuploidy and euploidy diploid cell classification. D: diploid; A: Aneuploidy. A-B: fibroblast cells from two patients (CRC15 was selected because only this sample Numbat could run successfully, other samples selected based on the largest number of autosomal aneuploidy cells in each cell type), C:T cells, D: B cells, E: endothelial cells.

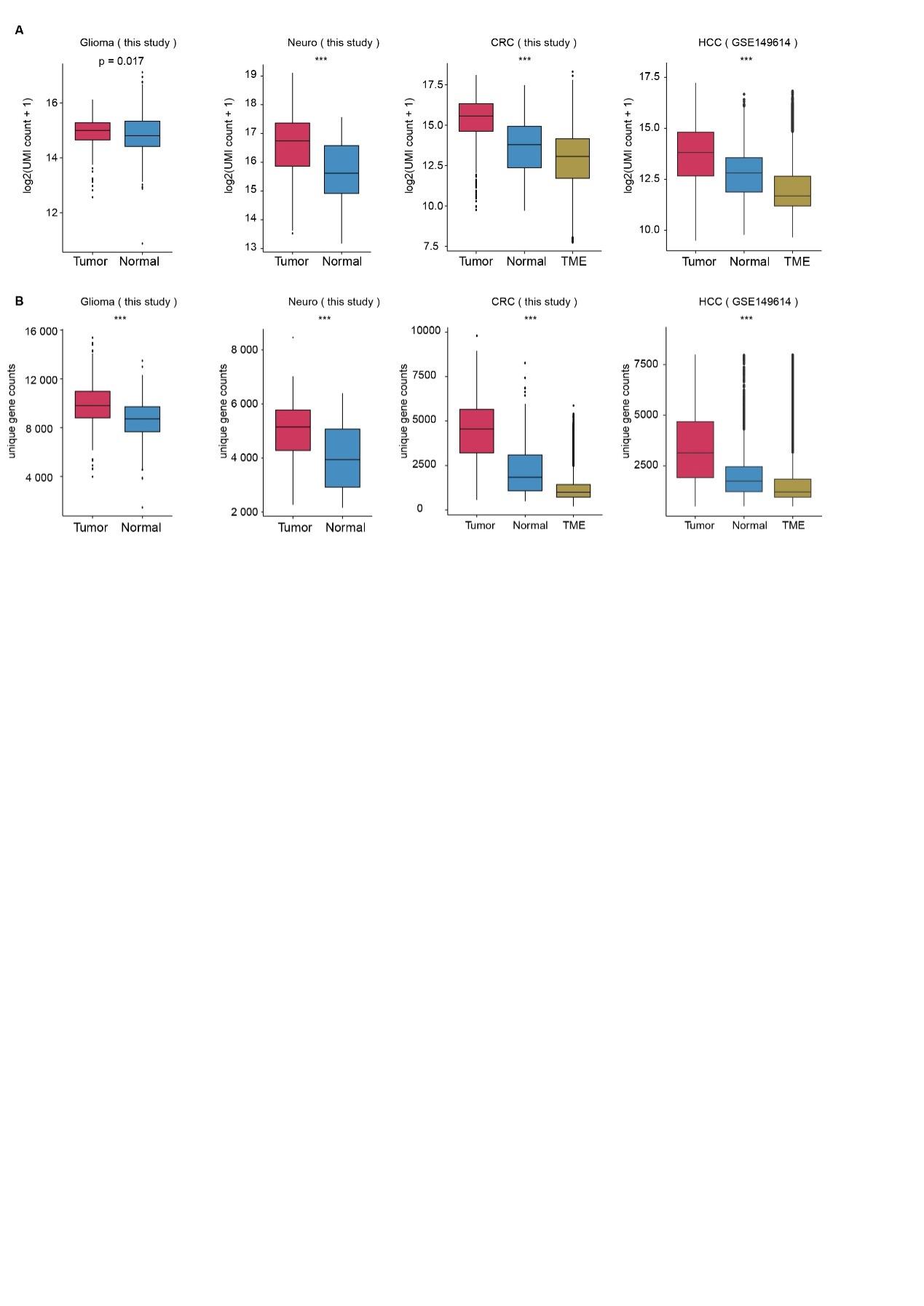

**Supplementary FigureS15:** **Distribution of cellular UMI and gene count in tumor and normal cells.**

A: Boxplot showing distribution of gene expression log2(UMI+1) in Tumor cells and normal cells

B: Boxplot showing distribution of unique gene counts in Tumor cells and normal cells

P values in A-Glioma, A-Neuro, B-Glioma and B-Neuro calculated using Wilcoxon rank-sum test and A-CRC, A-HCC, B-CRC and B-HCC calculated using Kruskal-Wallis test. NS: p > 0.05, *: p < 0.01, **: p < 0.001, ***: p < 0.0001

**Supplementary TableS1**. Benchmark datasets cells number

|  | Accession number |  | Tumor | Normal | TME |
| --- | --- | --- | --- | --- | --- |
| CRC | HRA000201 | CRC13 | 81 | 80 | 1088 |
|  |  | CRC11 | 42 | 113 | 1065 |
|  |  | CRC12 | 37 | 82 | 1058 |
|  |  | CRC16 | 29 | 49 | 508 |
|  |  | CRC02 | 65 | 9 | 431 |
|  |  | CRC22 | 58 | 6 | 551 |
|  |  | CRC15 | 4 | 57 | 317 |
|  |  | CRC03 | 41 | 20 | 622 |
| GBM | GSE185269 | Gliomas | 392 | 394 | 0 |
| Neuro | PRJCA002946 | Neuro | 5 | 37 | 0 |
| ALL | GSE144296 | ALL1 | 173 | 0 | 260 |
|  |  | ALL2 | 124 | 0 | 260 |
| NPC | GSE185269 | NPC43 | 137 | 0 | 0 |
| HUVEC | GSE185269 | HUVEC | 0 | 26 | 0 |

|  |  | Glioma | CRC13 | CRC11 | CRC12 | CRC16 | CRC02 | CRC22 | CRC15 | CRC03 | Neuro |
| --- | --- | --- | --- | --- | --- | --- | --- | --- | --- | --- | --- |
| N->N | scDNA | 394 | 80 | 113 | 82 | 49 | 9 | 6 | 57 | 20 | 37 |
|  | inferCNV | 393 | 80 | 111 | 80 | 49 | 0 | 6 | 57 | 20 | 37 |
|  | CopyKAT | 393 | 80 | 112 | 80 | 49 | 9 | 6 | 35 | 20 | 37 |
|  | SCEVAN | 393 | 0 | 0 | 0 | 0 | 0 | 0 | 0 | 0 | 37 |
|  | Numbat | 394 | 80 | 113 | 79 | 39 | 8 | 4 | 56 | 20 | 36 |
|  | CaSpER | 391 | 80 | 113 | 79 | 49 | 0 | 6 | 57 | 20 | 37 |
| T->N | scDNA | 0 | 0 | 0 | 0 | 0 | 0 | 0 | 0 | 0 | 0 |
|  | inferCNV | 17 | 0 | 0 | 0 | 1 | 65 | 31 | 0 | 21 | 0 |
|  | CopyKAT | 15 | 2 | 3 | 0 | 13 | 0 | 35 | 1 | 0 | 0 |
|  | SCEVAN | 16 | 0 | 0 | 0 | 0 | 0 | 0 | 0 | 0 | 0 |
|  | Numbat | 2 | 2 | 1 | 2 | 2 | 1 | 27 | 0 | 0 | 0 |
|  | CaSpER | 17 | 13 | 2 | 1 | 19 | 65 | 32 | 2 | 2 | 0 |
| N->T | scDNA | 0 | 0 | 0 | 0 | 0 | 0 | 0 | 0 | 0 | 0 |
|  | inferCNV | 1 | 0 | 2 | 2 | 0 | 9 | 0 | 0 | 0 | 0 |
|  | CopyKAT | 1 | 0 | 0 | 2 | 0 | 0 | 0 | 22 | 0 | 0 |
|  | SCEVAN | 1 | 69 | 105 | 79 | 48 | 9 | 5 | 52 | 20 | 0 |
|  | Numbat | 0 | 0 | 0 | 3 | 10 | 1 | 2 | 1 | 0 | 1 |
|  | CaSpER | 3 | 0 | 0 | 3 | 0 | 9 | 0 | 0 | 0 | 0 |
| T->T | scDNA | 392 | 81 | 42 | 37 | 29 | 65 | 58 | 4 | 41 | 5 |
|  | inferCNV | 375 | 81 | 42 | 37 | 28 | 0 | 27 | 4 | 20 | 5 |
|  | CopyKAT | 377 | 79 | 39 | 37 | 16 | 65 | 23 | 3 | 41 | 5 |
|  | SCEVAN | 376 | 80 | 41 | 37 | 28 | 65 | 56 | 4 | 41 | 5 |
|  | Numbat | 390 | 79 | 41 | 35 | 27 | 64 | 31 | 4 | 41 | 5 |
|  | CaSpER | 375 | 68 | 40 | 36 | 10 | 0 | 26 | 2 | 39 | 5 |

**Supplementary TableS2**. The true negative, false negative, false positive, and true positive in classification tumor vs normal cells.

**Supplementary Table S3**. Downsampling results of the average (each sequencing depth were sampled three times) sensitivity, specificity and accuracy for combined results from three samples

|  |  | Original | 10K | 3K | 1K |
| --- | --- | --- | --- | --- | --- |
| Sensitivity | inferCNV | 1 | 0.97 | 0.66 | 0.66 |
|  | CopyKAT | 0.97 | 0.95 | 0.80 | 0.78 |
|  | SCEVAN | 1 | 1 | 1 | 1 |
|  | Numbat | 0.97 | 0.53 | 0.29 | 0.27 |
|  | CaSpER | 0.92 | 0.71 | 0.58 | 0.41 |
| Specificity | inferCNV | 0.99 | 0.99 | 0.66 | 0.66 |
|  | CopyKAT | 0.99 | 1 | 1 | 0.88 |
|  | SCEVAN | 0 | 0 | 0 | 0 |
|  | Numbat | 0.99 | 0.74 | 0.60 | 0.70 |
|  | CaSpER | 0.99 | 0.99 | 1 | 0.98 |
| Accuracy | inferCNV | 0.99 | 0.98 | 0.66 | 0.66 |
|  | CopyKAT | 0.98 | 0.98 | 0.92 | 0.82 |
|  | SCEVAN | 0.38 | 0.45 | 0.54 | 0.71 |
|  | Numbat | 0.98 | 0.65 | 0.47 | 0.52 |
|  | CaSpER | 0.96 | 0.88 | 0.83 | 0.77 |

**Supplementary TableS4**. Computational speed time for each method.

| Sample | inferCNV | CopyKAT | SCEVAN | Numbat | CaSpER |
| --- | --- | --- | --- | --- | --- |
| CRC13 | CS: 5.12min | CS: 0.96min | CS: 0.79min | CS: 12.89min | CS: 2.75min |
| CRC11 | CS: 4.58min | CS: 0.96min | CS: 0.64min | CS: 12.24min | CS: 2.79min |
| CRC12 | CS: 3.51min | CS: 0.67min | CS: 0.55min | CS: 10.79min | CS: 2.02min |
| CRC16 | CS: 2.24min | CS: 0.51min | CS: 0.40min | CS: 7.68min | CS: 1.64min |
| CRC02 | CS: 1.81min | CS: 0.51min | CS: 0.43min | CS: 5.54min | CS: 1.52min |
| CRC22 | CS: 1.75min | CS: 0.37min | CS: 0.31min | CS: 5.35min | CS: 1.12min |
| CRC15 | CS: 1.28min | CS: 0.56min | CS: 0.34min | CS: 4.65min | CS: 1.02min |
| CRC03 | CS: 1.79min | CS: 0.44min | CS: 0.37min | CS: 5.39min | CS: 1.00min |

**Supplementary Note 1**. Selection of tools for benchmark.

| Tools | data requirement | years | citation | Lab(Last author) |
| --- | --- | --- | --- | --- |
| InferCNV | scRNA-Seq alone | NA | NA | NA |
| CONICSmat | scRNA-Seq+CNV regions | NA | NA | Aaron Diaz |
| Honeybadger | scRNA-Seq alone | NA | NA | Peter Kharchenko |
| Clonealign | scRNA-Seq+scDNA-Seq | NA | NA | Sohrab P. Shah |
| CopyKAT | scRNA-Seq alone | 2021 | 276 | Nicholas E. Navin |
| SCEVAN | scRNA-Seq alone | 2023 | 9 | Michele Ceccarelli |
| Numbat | scRNA-Seq alone | 2022 | 25 | Peter Kharchenko |
| CaSpER | scRNA-Seq alone | 2020 | 107 | Xiaobo Zhou |

We collected the tools information at 2023(See Table above). We first collected all the tools available that only required scRNA-Seq data without any other data requirement. For example, CONICSmat was not included as it required users to specify CNV regions to run. Clonealign was not included as it required both scRNA-Seq and scDNA-Seq from the same tumor. Next, we filtered out tools that were not used widely (fewer than 20 citation per year for methods published more than two years). Lastly, if multiple tools were developed from the same lab and the publication already suggested that the newer method was superior than the older method, only the newer method would be kept. For example, the Honeybadger was not included due to this reason (Numbat developed from the same lab and was shown to perform better than Honeybadger).

**Supplementary Note 2**. Selection of tools for breakpoint evaluation.

Because of the relative low sequencing depth of scDNA-Seq, it is difficult to obtain an accurate single cell level breakpoint detection and evaluation. InferCNV, Numbat and SCEVAN reported the clonal breakpoints, therefore we evaluated the breakpoints for these three tools..

Although CopyKAT could generate single cell level segments files, but no clear instructions were given on how to obtain a clonal breakpoint calling. Previously, in Numbat manuscript (*Haplotype-aware analysis of somatic copy number variations from single-cell transcriptomes*, **Nature Biotechnology**, 2022), they used a <-0.03 cutoff to define copy number deletion and a >0.03 cutoff to define copy number amplification for CopyKAT. We’ve looked carefully into the data, we were uncertain about where to draw the line as the cutoff. And such cutoff significantly impacts the final results. CopyKAT manuscript itself also didn’t provide a recommended cutoff. CaSpER does not include the breakpoint detection functionality, thus is not included in our breakpoint evaluation.

**Supplementary Note 3**. Reference cell selection for CNV inference for tumor.

Among the five tools we tested, the input of reference cells is required as input for Numbat and CaSpER and optional for inferCNV, CopyKAT and SCEVAN. Notably, CopyKAT and SCEVAN developed built-in function to optimize the selection of reference cells. CopyKAT utilizes a combined approach to automatically identify high-confidence normal cells for reference, while SCEVAN leverages predefined normal cell gene signatures, including those from stromal and immune cells, to identify high-confidence normal cells. Therefore, in most cases, you do not need to optimize the reference cells for CopyKAT and SCEVAN. But it is recommended to optimize the reference cell selection for other methods.

When selecting high-confidence normal cells, we recommend adhering to several key principles to ensure robust CNA profile calculations.

(1) it is best to choose same cell types for your reference (normal) and observation (tumor) from the same dataset. This is based on recommendation from almost every tools, which can be found both in the reference publication as well as the developer’s replies on the Github issues. For example, if your tumor cells are from epithelial tissues, then it will be better to use nonmalignant epithelial cells from the same tissue as reference cells. This approach minimizes cell-specific expression patterns and differences, ensuring that the expressed genes are consistent across both groups. Some of our preliminary testing also supported this recommendation (See Figure R1 below: CNV profiles estimated from CopyKAT showed that using normal alone without TME cells are more similar to the DNA CNA ground truth).

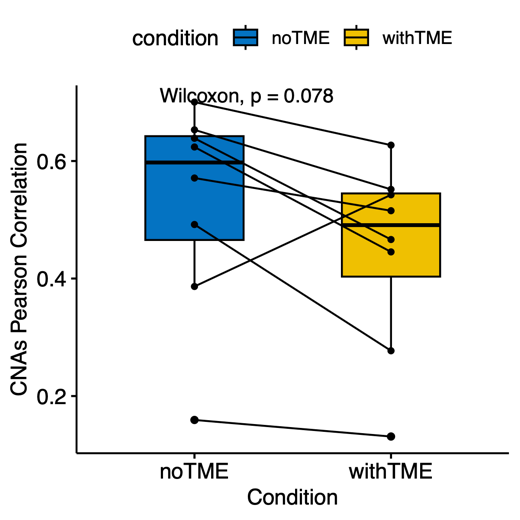

Figure R1: CNA profile performance of CopyKAT with or without TME cells

Therefore, it is crucial to identify the normal reference cells from your input scRNA-Seq dataset. Let’s take CRC datasets as an example, first we will obtain the epithelial cluster from other tumor microenvironment cell types. Then epithelial cluster can be further separated into more clusters. There are few ways you can identify the nonmalignant cells, (a) we found in most cases nonmalignant cells tend to have a lower UMI count compared with malignant cells, which is due to the common hypertranscription in cancer (*Widespread hypertranscription in aggressive human cancers*, **Science Advances**, 2022). So you can find the cluster with the lowest total UMI counts; (b) To annotate reference (normal) cell types, you can try to assess marker genes expression (for example, Numbat used high expression of EPCAM gene to identify breast cancer tumor cells from normal cells, and higher expression of KRT8 to identify anaplastic thyroid cancer cells. This has to be based on literature and users own data. Therefore, both approaches can be combined to identify correct normal cells (cell type of tumor origin). In addition, if normal cells were used incorrectly, you might detect the opposite CNA profiles. This can be identified by checking whether the detected CNA is consistent with known frequent large CNA events for that tumor type. For example, 8q is normally observed as a copy number gain in tumor, if you detected copy number loss, it may indicate that the wrong reference cells were selected.

(2) if there are not sufficient normal cells, you can choose either to (a) add other cell types, such as TME cells (fibroblast, endothelial, immune cells) from the same dataset (Figure 2E-F) or (b) add external expression matrix as suggested by Numbat. For example, Numbat itself indeed has a build-in expression matrix which can be used as a reference.

To summarize, we’ve also presented a decision tree (Figure R2) to facilitate the users to select proper reference cells.

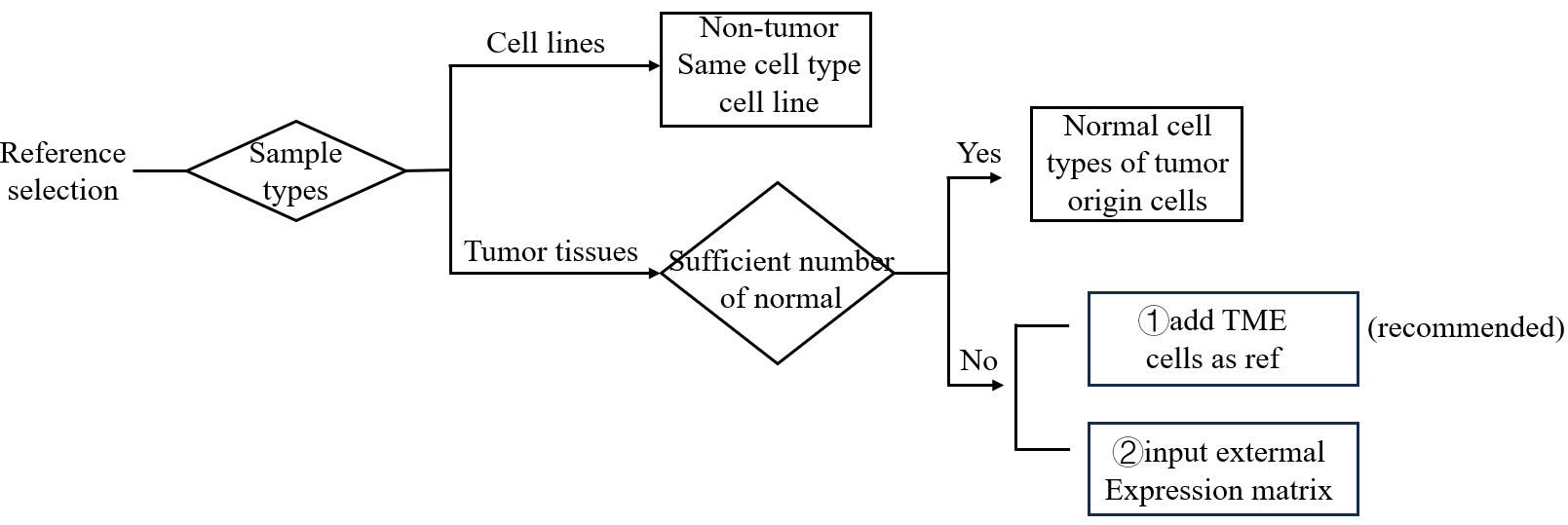

Figure R2: the decision tree for reference cell selection
